# The metastable brain associated with autistic-like traits of typically developing individuals

**DOI:** 10.1101/855502

**Authors:** Takumi Sase, Keiichi Kitajo

**Affiliations:** Rhythm-based Brain Information Processing Unit, CBS-TOYOTA Collaboration Center, RIKEN Center for Brain Science, Wako, Saitama 351-0198, Japan; Department of Electrical Engineering, Faculty of Engineering, University of Malaya, Kuala Lumpur 50603, Malaysia; Division of Neural Dynamics, Department of System Neuroscience, National Institute for Physiological Sciences, National Institutes of Natural Sciences, Okazaki, Aichi 444-8585, Japan; Department of Physiological Sciences, School of Life Science, The Graduate University for Advanced Studies (SOKENDAI), Okazaki, Aichi 444-8585, Japan

## Abstract

Metastability in the brain is thought to be a mechanism involved in dynamic organization of cognitive and behavioral functions across multiple spatiotemporal scales. However, it is not clear how such organization is realized in underlying neural oscillations in a high-dimensional state space. It was shown that macroscopic oscillations often form phase-phase coupling (PPC) and phase-amplitude coupling (PAC) which result in synchronization and amplitude modulation, respectively, even without external stimuli. These oscillations can also make spontaneous transitions across synchronous states at rest. Using resting-state electroencephalographic signals and the autism-spectrum quotient scores acquired from healthy humans, we show experimental evidence that the PAC combined with PPC allows amplitude modulation to be transient, and that the metastable dynamics with this transient modulation is associated with autistic-like traits. In individuals with a longer attention span, such dynamics tended to show fewer transitions between states by forming delta-alpha PAC. We identified these states as two-dimensional metastable states that could share consistent patterns across individuals. Our findings suggest that the human brain dynamically organizes inter-individual differences in a hierarchy of macroscopic oscillations with multiple timescales by utilizing metastability.

**Author Summary:** The human brain organizes cognitive and behavioral functions dynamically. For decades, the dynamic organization of underlying neural oscillations has been a fundamental topic in neuroscience research. Even without external stimuli, macroscopic oscillations often form phase-phase coupling and phase-amplitude coupling (PAC) that result in synchronization and amplitude modulation, respectively, and can make spontaneous transitions across synchronous states at rest. Using resting-state electroencephalography signals acquired from healthy humans, we show evidence that these two neural couplings enable amplitude modulation to be transient, and that this transient modulation can be viewed as the transition among oscillatory states with different PAC strengths. We also demonstrate that such transition dynamics are associated with the ability to maintain attention to detail and to switch attention, as measured by autism-spectrum quotient scores. These individual dynamics were visualized as a trajectory among states with attracting tendencies, and involved consistent brain states across individuals. Our findings have significant implications for unraveling variability in the individual brains showing typical and atypical development.

## Introduction

The human brain can spontaneously yield transition dynamics across oscillatory states and organize a variety of events in a hierarchy of oscillations. Such spontaneous dynamics, particularly at rest, have been intensively observed and analyzed over many years, and attempts to model them have been made using dynamical systems theory, to achieve a better prediction of brain activity [1–5]. However, there is a lack of direct evidence that resting-state brain dynamics can originate from the underlying nonlinear states, and little is known about the kind of states that may have functional roles in the dynamic organization of spontaneous activity in a way utilizing oscillatory hierarchy.

Electroencephalography (EEG) is a promising technique for the high temporal resolution observation of the dynamics of neural activity over large-scale brain networks. The observed macroscopic signals are oscillatory such that the corresponding power spectrum exhibits a representative peak, and are classified into multiple bands according to its frequency in general [6, 7]. The peak frequency of neural activity shows either a higher or lower value depending on brain function [6] and cognitive and behavioral performance [7]; for example, alpha-band activity can be enhanced or suppressed by attention, and its peak frequency can vary with age [7].

Observed macroscopic neural oscillations reflect underlying nonlinear dynamics. Experimental studies have presented evidence that phases detected from oscillations at a particular frequency show the nonlinear brain phenomenon called phase synchronization by forming phase-phase coupling (PPC) [8–10]. Furthermore, the phases have the ability to modulate the amplitude of a faster oscillatory component by forming phase-amplitude coupling (PAC) [11–15]; in recent years, PAC was observed not only locally but also between regions of the large-scale network [16, 17]. The PPC has been suggested to play a role in making functional connections among distant brain regions [8], while it is suggested that the PAC mediates computation between local and global networks [12], with both couplings having been observed in function-specific and individual behavior-related oscillations at multiple spatiotemporal scales [9, 14]. From a dynamical systems theory point of view, these two kinds of coupled oscillatory dynamics can be interpreted as being generated from coupled oscillatory attractors composed of the limit cycle [2] or its variant form, i.e., a torus in a high-dimensional state space [18, 19]. These suggestions have inspired phenomenological modeling of the dynamics underlying EEG neural oscillations [20, 21], resulting in a variety of coupled nonlinear-oscillator systems as represented by the Kuramoto model [20–23].

Oscillatory dynamics such as those mentioned above can make spontaneous transitions among multiple network states, particularly at rest. Previous studies labeled EEG signals observed during a resting condition as a small number of states called microstates [24–27]. These microstates have been suggested to be associated with cognition and perception [25], as well as individual differences in brain function [27]. In recent years, resting-state EEG signals have been investigated from the point of view of functional connectivity of the large-scale network, which is often characterized by the strength of the PPC [28–30]. Betzel et al. showed evidence that resting-state EEG phases exhibit dynamic changes in PPC modulation so that a repertoire of synchronized states appears [29]. Moreover, PAC modulation from EEG phases to amplitudes at rest was recently studied [31, 32] because the PAC can also occur spontaneously [13]; inter-regional PAC during rest was also intensively analyzed [33]. These experimental findings imply that resting-state EEG phase dynamics not only exhibit synchronization but also result in amplitude modulation over the large-scale network at the same time via both PPC and PAC. Therefore, we developed the following hypotheses for the resting brain: (i) there is a repertoire of synchronous slow oscillations that interact via the PPC and interact with fast oscillations via the PAC (Fig 1A); (ii) these synchrony-dependent slow and fast oscillations result in a repertoire of PAC states characterized by slow and fast timescales (Fig 1B); and (iii) transitions between metastable PAC states, i.e., dynamic and large-scale changes in PAC strengths, occur spontaneously at rest according to transitions among a repertoire of the synchronous slow oscillations, that is, according to the dynamic changes in PPC strengths (Fig 1C).

**Fig 1.**
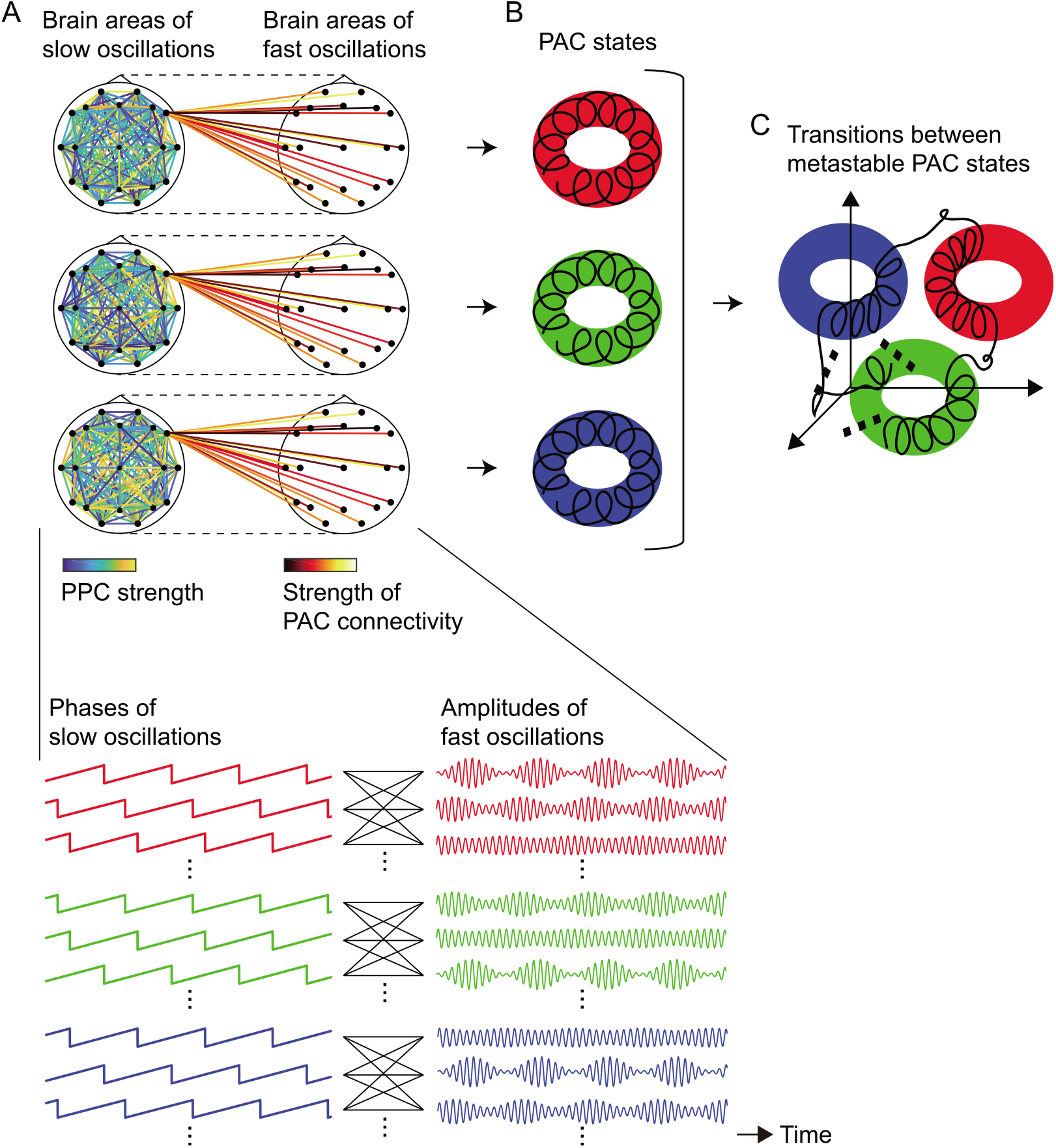
The dynamic PPC-PAC hypothesis. (A) A repertoire of synchronous slow oscillations that interact via the PPC and interact with fast oscillations via the PAC. (B) The resulting PAC states. (C) Transitions between the metastable PAC states. The dynamic PPC-PAC hypothesis states that for the resting brain, dynamic changes in PPC strengths (transitions between synchronous states) can cause dynamic and large-scale changes in PAC strengths because of PPC-PAC connectivity (A), and thereby yield transitions between oscillatory states with multiple peak frequencies (B and C). The oscillations of each state can realize the transition to another state by spontaneous fluctuations in the brain; in other words, the underlying states can show metastability.

In this study, we aimed to test the dynamic PPC-PAC hypotheses described above (Fig 1), and to show experimental evidence of how the metastable human brain is associated with autistic-like traits. In recent years, metastability in the brain has been proposed as a mechanism for integration and segregation across multiple levels of brain functions [34]. To elucidate aspects of the dynamics of the metastable human brain in this study, we first developed a method to label observed metastable dynamics as the underlying states in a data-driven manner based on the oscillatory hierarchy hypothesis [35]. The method was then applied to 63-channel high-density scalp-recorded EEG signals from 130 healthy humans in an eyes-closed resting condition (*n* = 162 in total; 32 subjects participated in the experiment twice). The obtained results were compared with the autism-spectrum quotient (AQ) subscales [36, 37] acquired from 88 of the subjects after the EEG recording, and were combined with the model of a coupled oscillator system driven by spontaneous fluctuations to validate our hypothesis (Fig 1).

## Materials and Methods

### Data Acquisition

In total, 130 healthy humans (Age: 24.0 ± 5.0 years, mean ± SD, 66 females) participated in the EEG experiment after giving written informed consent. The EEG study was approved by the ethics committee of RIKEN (the approval number: Wako3 26-24) and was conducted in accordance with the code of ethics of the Declaration of Helsinki. Thirty-two subjects participated in the experiment twice. The EEG signals were recorded from an EEG amplifier (BrainAmp MR+, Brain Products GmbH, Gilching, Germany) and a 63-channel EEG cap (Easycap, EASYCAP GmbH, Herrsching, Germany) placed on the scalp in accordance with the International 10/10 system with a left earlobe reference and AFz as a ground electrode. The signals were recorded for 180 s with the subjects in an eyes-closed resting condition. The following experimental configuration was used: sampling frequency 1000 Hz, low-cut frequency 0.016 Hz, and high-cut frequency 250 Hz. The recorded signals were offline re-referenced to the average potentials of the left and right earlobes. The EEG data were also analyzed in our previous studies [38, 39], which had a different purpose. After the EEG experiment, 88 subjects were asked to answer the Japanese version of the AQ questionnaire [37], which was originally constructed by Baron-Cohen et al. (2001) [36]. The following five AQ subscales were scored from the obtained answers: social skills, attention to detail, attention switching, communication, and imagination. All the analyses shown below were conducted using in-house code custom written in MATLAB (Mathworks, Natick, MA, USA) with the EEGLAB [40], FieldTrip [41], and CSD Toolboxes [42]. The code [43–45] was also used after minor changes.

### Metastable States Clustering

Metastable states clustering was developed in a data-driven manner to label observed metastable dynamics as the underlying states. The method consisted of the following three analyses: (i) multi-step envelope analysis, (ii) k-means clustering, and (iii) supervised dimensionality reduction of state space by linear discriminant analysis (LDA) [46] (see Fig 2). Analysis (i) was motivated by the Poincaré section in flows and sections in maps [47] with their application to a modeled neural network [18]; these sections enabled to convert a *d*-dimensional torus in a state space to the (*d* − 1)-dimensional torus recursively [47]. We incorporated this idea into signal processing of resting-state EEG data, and aimed to transform the original transition dynamics (Fig 2A) into a trajectory among zero-dimensional states (Fig 2D), to which the potential energy can be applied. In this study, we realized this multi-step conversion by repeatedly computing envelopes from EEG signals (Fig 2B). A vast amount of studies used such envelope analysis so that the nested oscillations (with slow and fast timescales) changed to amplitude oscillations (with a slow timescale) [13, 48].

**Fig 2.**
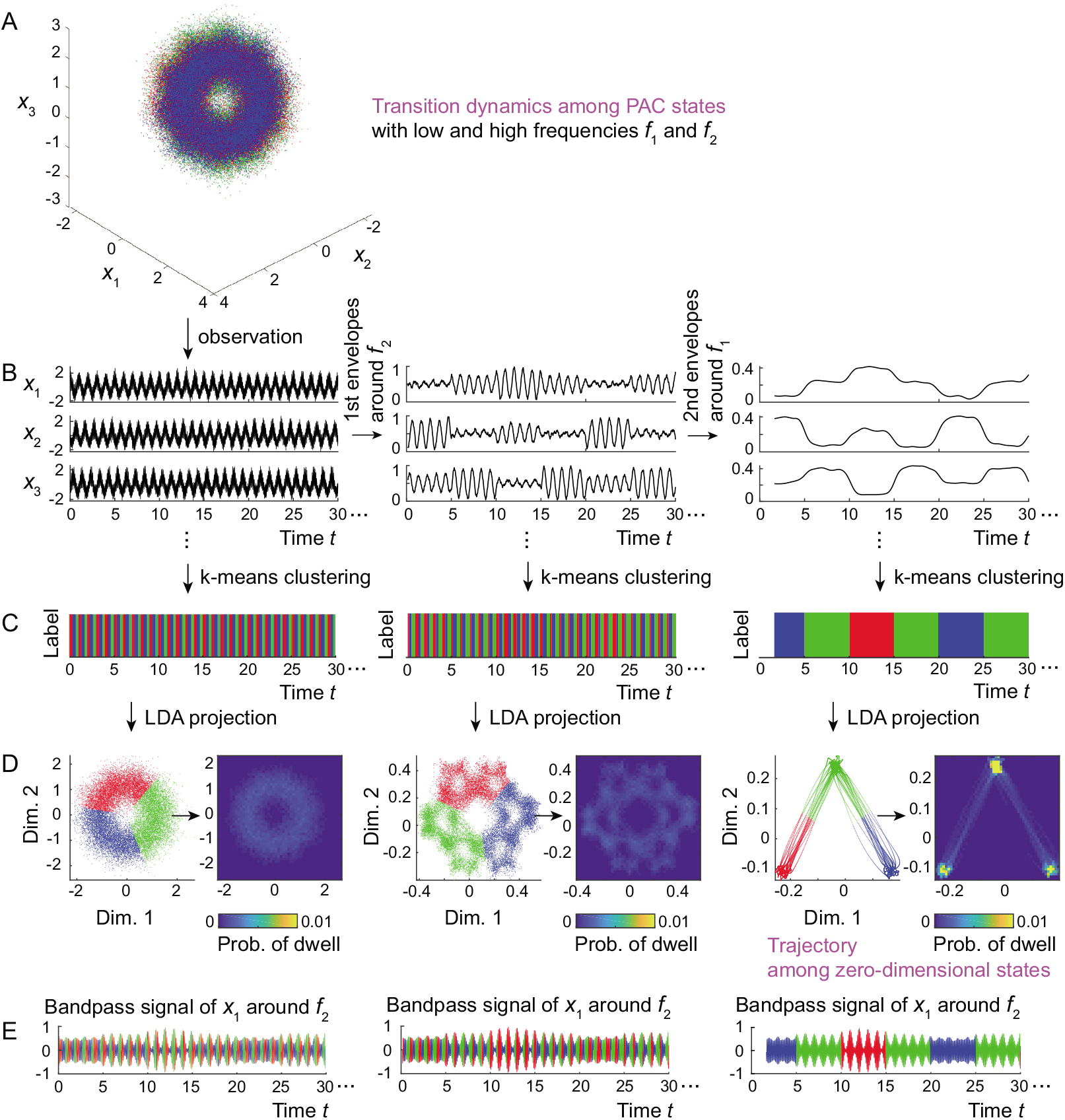
The flow of signal processing in metastable states clustering. (A) A transition dynamics among three PAC states (*d* = 2) in the state space of *x*_*i*_(*t*) = cos(2*πf*_1_*t* + *ψ*_*i*_) + 0.5[1 + *b*_*i*_(*t*) cos(2*πf*_1_*t* + *ψ*_*i*_)] sin(2*πf*_2_*t*) + *ξ*_*i*_(*t*), where the first, second, and last terms indicated the slow oscillations, amplitude-modulated fast oscillations, and noise, respectively. (B) Observed signals, the first envelopes (instantaneous amplitudes) around *f*_2_, and the second envelopes around *f*_1_. (C) The labeled sequences. (D) LDA projections of the state space. (E) The bandpass signal of *x*_1_(*t*) around peak frequency *f*_2_ and corresponding labels. The frequencies *f*_1_ and *f*_2_ here were set to 1 Hz and 10 Hz, respectively; modulation index *b*_*i*_(*t*) dynamically changed among strengths 0.15, 0.5, and 0.85; and noise *ξ*_*i*_(*t*) followed a normal distribution of mean 0 and standard deviation 0.3. The colors in panel A correspond to those in the right column of panels C to E.

The method was applied to the resting-state scalp EEG data with the dimension *N* = 63, under the assumption that (I) the underlying system generates the dynamics of transitions between states with *d* frequencies *f*_1_, *f*_2_, .., *f*_*d*_ (changes in strengths of hierarchical coupling), such that slow oscillations with *f*_*i*_ hierarchically modulate fast ones with *f*_*i*+1_ for *i* = 1, 2, …, *d* − 1 (see [35] for the oscillatory hierarchy hypothesis). Moreover, we assumed that (II) the observation process is the linear sum of distinct oscillations. These assumptions allowed the underlying states to be *d*-dimensional, and the *N* recorded EEG signals to reflect the common neural oscillations with *d* peak frequencies. Hereafter, we denoted these *N* signals by ***X***(*t*) = (*X*_1_ (*t*), *X*_2_ (*t*), …, *X*_*N*_ (*t*)) for *t* = 0 to 180 s (:=*T*). As shown in the detailed signal processing steps below, analyses (i) to (iii) tested whether the resulting states can be seen as zero-dimensional states with attracting tendencies (which we called the weakly attracting states) after *d*-time computations of envelopes. In other words, we tested whether the states of the original transition dynamics can be *d*-dimensional metastable states. Figure 2 illustrates that the PAC states (Fig 2A) are identified as the two-dimensional metastable states through transformation into the zero-dimensional weakly attracting states (in Fig 2D), whose labels clearly reflect different strengths of amplitude modulation (in Fig 2E).

Analysis (i): First, the instantaneous amplitudes were repeatedly computed from signals ***X*** (*t*) around frequencies *f*_*d*_, *f*_*d*−1_, …, *f*_1_, as follows (Fig 2B):

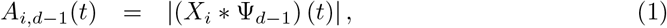

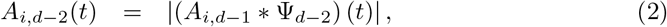

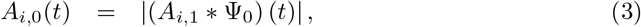

for *i* = 1, 2, …, *N*, where Ψ_*d*−1_ (*t*), Ψ_*d*−2_ (*t*), …, Ψ_0_ (*t*) were the complex-valued Morlet wavelets [49, 50] defined as

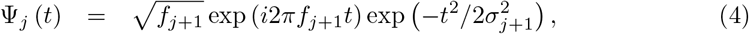

for *j* = 0, 1, …, *d* – 1. Operators (·∗·) and |·| denote a convolution and conversion from the complex value to its amplitude, respectively, and we defined ***A***_*j*_(*t*) = (*A*_1,*j*_(*t*), *A*_2,*j*_(*t*), …, *A*_*N,j*_(*t*)). To obtain the results at high temporal resolution, we set *σ*_*j*+1_ to a value such that the number of cycles *n*_co_ of the wavelet Ψ_*j*_ (*t*) was three, i.e., *n*_co_:=6*f*_*i*_*σ*_*j*+1_ = 3. The data ***A***_*j*_(*t*) for *t* = *n*_co_/2*f*_*j*+1_ to (*T* − *n*_co_/2*f*_*j*+1_) were used to reduce the edge artifact of the wavelet Ψ_*j*_ (*t*) with update *T* − *n*_co_/*f*_*j*+1_ *→ T*, with respect to each *j*. The peak frequency *f*_*d*_ and others *f*_*d*−1_, *f*_*d*−2_, …, *f*_1_ were estimated from the power spectra of *X*_*i*_(*t*) and of *A*_*i,d*−1_(*t*), *A*_*i,d*−2_(*t*), …, *A*_*i*,1_(*t*), respectively, for *i* = 1, 2, …, *N*. We averaged each set of the spectra over *i* with respect to each frequency *f*, obtained a single spectrum, and estimated its peak frequency over the interval 1≤*f* < 45 for ***X***(*t*), otherwise in 0.1≤*f* < *f*_*j*+1_ for ***A***_*j*_(*t*); for ***X***(*t*) only, we first reduced the power-law effect on the corresponding spectra *P*_*i*_(*f*) across *i* = 1, 2, …, *N* which may follow *f* ^−*β*^1, *f* ^−*β*^2, …, *f* ^−*β*^*N* with certain exponents [51], respectively, by subtracting the linear trend from log *P*_*i*_(*f*) vs. log *f* with respect to each *i*.

Analysis (ii): Next, instantaneous amplitudes ***A***_0_(*t*) were labeled as the *K* states of *L* (*t*) *∈{*1, 2, …, *K}* by k-means clustering (Fig 2C). The number of states *K* was estimated from the Calinski-Harabasz index [52] in a condition of *K* ∈{\ 2, 3, …, 10}. To obtain reproducible results, we deterministically initialized the clustering algorithm using PCA partitioning [53]. Any kernel function was not applied to the present clustering analysis because the dynamics ***A***_0_ (*t*) appeared here would play a role representing *K* states more discriminatorily in the *N*-dimensional state space compared with original dynamics ***X*** (*t*) (see Figs 2A and 2D).

Analysis (iii): Finally, the labeled amplitudes (***A***_0_(*t*), *L*(*t*)) were projected to lower-dimensional data ***Y*** (*t*) = (*Y*_1_(*t*), *Y*_2_(*t*), …, *Y*_*m*_(*t*)) with dimension *m < N* by LDA (Fig 2D). In this study, (***A***_0_(*t*), *L*(*t*)) was projected onto a plane (*m* = 2) for the case of *K* > 2, otherwise in a one-dimensional axis (*m* = 1) due to limitation of the LDA. We generated histograms with respect to each labeled state *k* ∈ {1, 2, …, *K*} using the same bin sizes, and calculated the maxima of the counts of bins *E*_*k*_ for each *k*. The statistics {*E*_*k*_} can indicate higher values as states {1, 2, …,*K*} become stable (see Fig 2D), and were regarded as the indices for the attracting tendencies that could be proportional to the potential energy of state space. The statistic *E* = min_*k*_ *E*_*k*_ was applied to the Fourier-transform (FT) surrogate data testing for multivariate time series [54] under the null hypothesis *H*_0_, where ***A***_0_ (*t*) is linearly correlated Gaussian noise. We generated surrogate data 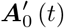 by randomizing the initial phases of ***A***_0_ (*t*), applied k-means clustering 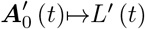, converted the labeled data 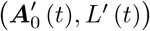 to lower-dimensional ones ***Y′*** (*t*) by the same projection as (***A***_0_ (*t*), *L* (*t*)), and calculated the statistic *E′* of the surrogate data. We performed a one-sided test to verify whether *E* was significantly larger than *E′* by generating 200 surrogate data sets and setting the significance level to 0.05.

In summary, ***X*** (*t*) was converted into ***Y*** (*t*) using the following composite analysis: (i) the envelopes were repeatedly computed from 63-channel EEG signals *d* times (Fig 2B); (ii) the final envelopes were labeled as *K* clusters in the 63-dimensional space (Fig 2C); and (iii) these clusters were projected onto a two- or one-dimensional space in which we tested whether they showed attracting tendencies (Fig 2D). These analyses were performed in the conditions of *d ∈ {*0, 1, 2, 3}, and we tested which *d* can indicate the dimension of underlying states. For the case of *d* = 0 only, we first applied a band-pass filter to the raw EEG signals in a range between 1 and 45 Hz.

### Dynamical Systems Modeling

We developed a model for individual transition dynamics, in which the dimension of underlying states was identified as two. The model was described by a coupled oscillator system driven by fluctuations, and was combined with the empirical results to validate the dynamic PPC-PAC hypothesis (Fig 1). The large-scale network with PPC-PAC connectivity was modeled in a space spanned by the 63 electrodes, and each node was represented by the phase of slow oscillations and the amplitude of fast oscillations (i.e., the PAC oscillator; S1 Fig). Inter-individual differences in the model were then taken into account through the PPC connectivity (see Fig 1A), which was estimated from the current source density (CSD) [42, 55] of the raw EEG signals ***X***(*t*). Finally, we applied an observation operator *h* from CSD to EEG dynamics.

The signals ***X*** (*t*) (represented as the column vector hereafter) were applied to two (*N* × *N*)-matrices ***G*** and ***H*** and were transformed into CSD signals ***I***_m_ (*t*) as follows [42, 55, 56]:

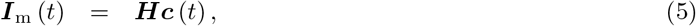

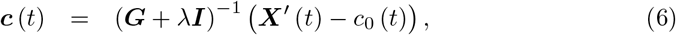

where

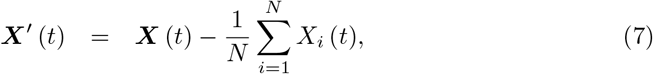

and the following spherical interpolation was used [55]:

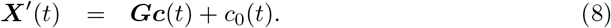

In Eq (6), ***c***(*t*) and *c*_0_(*t*) are the spline coefficients, *λ* denotes the smoothing constant which was set to 10^−5^, and ***I*** denotes the identity matrix with size *N*. As seen in Eqs (5) to (8), ***I***_m_(*t*) can be mapped to ***X***(*t*) through observation *h* as follows:

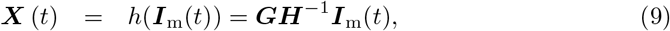

under the assumption that *c*_0_ (*t*) and the average potential in Eq (7) are zero. As shown in the Results, many of the individual transition dynamics were labeled as two-dimensional metastable states with delta-alpha PAC, and thus a coupled delta-alpha PAC oscillator system was constructed.

### The Model

The delta-alpha PAC dynamics of ***I***_m_(*t*) were modeled by a coupled oscillator system driven by fluctuations (Fig S1). The model comprised *N* PAC oscillators whose phases and amplitudes corresponded to delta-(*δ*) and alpha-band (*α*) activity, respectively. We represented the *i*th oscillator at time *t* by delta-band phase *ϕ*_*i*_ (*t*) and alpha-band amplitude *r*_*i*_ (*t*), its frequency by *ω*, and the fluctuation to this oscillator by *η*_*i*_ (*t*). The phases *ϕ*_*j*_ (*t*) have connections to *ϕ*_*i*_ (*t*) (PPC) for *j≠i* with coupling strengths 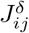 and delay 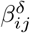, and connections to amplitude *r*_*i*_ (*t*) for both *j* = *i* and *j≠i* (local and inter-regional PAC) with coupling strengths 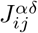 and delay 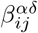. The phase *ϕ*_*i*_ (*t*) has another connection from fluctuation *η*_*i*_ (*t*) with the level *D*. Then, the system with these three kinds of interactions can be described as follows:

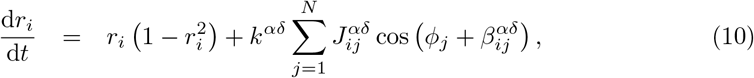

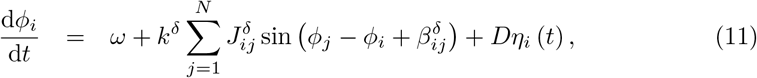

where *k*^*δ*^ and *k*^*αδ*^ denote the total coupling strength of the PPC and PAC, respectively. In Eq (11), term 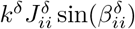 (where *j* = *i* in summation) was assumed as a certain bias arising from the estimation of PPC connectivity, shown below. It can be seen that a part of the system composed only of delta-band phases (Eq (11)) is the Kuramoto model (or also called the Kuramoto-Sakaguchi model) subject to noise [22]. Moreover, the present system (Eqs (10) and (11)) can reduce to the identical Stuart-Landau oscillators when *k*^*δ*^, *k*^*αδ*^, and *D* are all set to zero [22]. In this model, we made a connection from fluctuation *η*_*i*_(*t*) to phase *ϕ*_*i*_(*t*) only because our hypothesis (Fig 1) was that the dynamic changes in PAC strengths (transitions between PAC states) can be attributed to changes in PPC strengths (transitions between synchronous states). Accordingly, we set 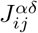 and 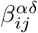 in Eq (10) to arbitrary values that could not affect the transition dynamics.

As described below, synchronous states in the model were represented by phase patterns with different relative phase relationships, and were thus characterized by a set of the phase differences [57]. The model (Eqs (10) and (11)) without the fluctuation term *Dη*_*i*_(*t*) can show multistability where each attractor generates oscillations with a distinct phase pattern, and such oscillations in a state can realize the transition to another state by fluctuations *Dη*_*i*_(*t*) (see the scenario of multistability in [58]). In the following, the delta-band phase patterns were first estimated from the individual CSD signals ***I***_m_ (*t*). The delta-band PPC connectivity 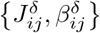 and the level of fluctuations *D* were then estimated in turn. Finally, the model Eqs (10) and (11) was numerically simulated with *ω* = 2*πf*_1_, *k*^*δ*^ = *k*^*αδ*^ = 0.1 and ***η***(*t*) = (*η*_1_(*t*), *η*_2_(*t*), …, *η*_*N*_ (*t*)) being generated from a multivariate normal distribution, using the Euler-Maruyama method [59] with time step Δ*t* = 0.01 s. Hereafter, we represented the PPC-PAC connectivity by complex-valued matrices ***C***^*δ*^ and ***C***^*αδ*^, which were defined as 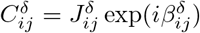 and 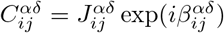, respectively.

### Estimation of Phase Patterns

Phase patterns of the delta-band oscillatory dynamics were estimated from CSD signals ***I***_m_ (*t*) based on the label *L* (*t*) ∈ {1, 2, …, *K}* of ***A***_0_(*t*). First, delta-band phases ***θ***(*t*) = (*θ*_1_(*t*), *θ*_2_(*t*), …, *θ*_*N*_ (*t*)) were extracted from ***I***_m_ (*t*) using the complex-valued Morlet wavelet Ψ_0_ (*t*) defined by Eq (4), as follows:

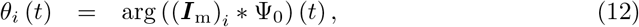

where arg (*·*) denotes the conversion from the complex value to its phase. The parameter *σ*_1_ of Ψ_0_(*t*) in Eq (4) was set to a value, such that *n*_co_ = 3 as well as Ψ_0_(*t*) applied to ***A***_1_(*t*). Next, the time courses of the phase differences between each pair of *θ*_*i*_ (*t*) and *θ*_*j*_ (*t*) were calculated, and were averaged over time with respect to each labeled state *µ* as follows:

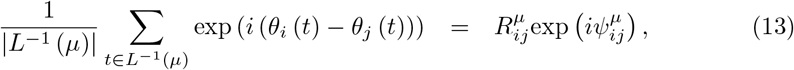

where |*·*| denotes the number of time instances at which the dynamics visited state *µ*. The absolute and argument parts 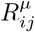 and 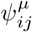 in Eq (13) indicated the mean of the phase synchronization index [49] and the mean of the phase difference between *θ*_*i*_ (*t*) and *θ*_*j*_ (*t*) in state *µ*, respectively. Then, the phase pattern (the relative phases) 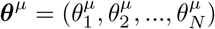 was estimated from phase differences 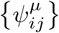 according to the following algorithm composed of initialization steps

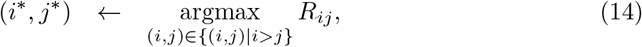

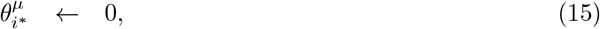

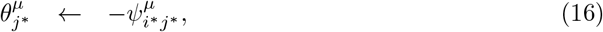

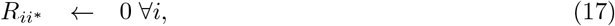

and (*N* − 2)-time iterations of

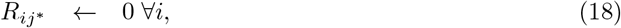

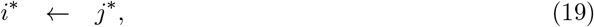

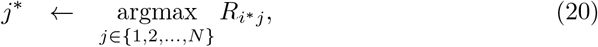

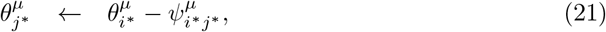

for *µ* = 1, 2, …, *K*, where the following index has been defined:

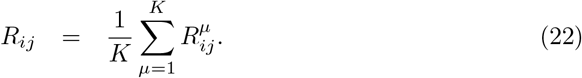

The index *R*_*ij*_ was used to measure how reliably the phases *θ*_*i*_ (*t*) and *θ*_*j*_ (*t*) were phase-locked over time and among states. As a whole, the phase pattern ***θ***^*µ*^ was regarded as the synchronous state *µ* in the model.

### Estimation of PPC Connectivity

The PPC connectivity ***C***^*δ*^ was estimated from synchronous states {***θ***^*µ*^} as follows [57]:

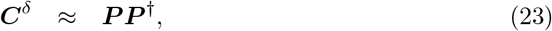

where

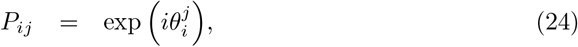

and ***P***^†^ denotes the Moore-Penrose inverse of ***P***. The obtained connectivity ***C***^*δ*^ can converge a trajectory of the system Eq (11) into either of synchronous states {***θ***^*µ*^} such that 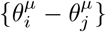 are preserved [57].

### Estimation of the Level of Fluctuations

The fluctuation level *D* was estimated from system Eq (10) with PPC connectivity ***C***^*δ*^. The fluctuations ***η***(*t*) were generated from the multivariate normal distribution of mean zeros and the covariance matrix, which was represented by a nearest symmetric positive semidefinite matrix [60] of matrix **Σ** defined as

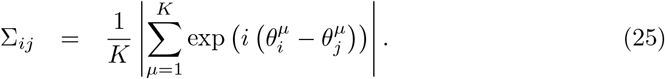

The value Σ_*ij*_ measures the similarity of 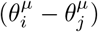 among synchronous states {***θ***^*µ*^}; as 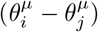 differs, Σ_*ij*_ approaches zero, and so the fluctuations *η*_*i*_(*t*) and *η*_*j*_(*t*) become independent of each other. That is, we assumed that the delta-band phase difference (*θ* _*i*_ (*t*) − *θ*_*j*_(*t*)) was likely to fluctuate as it showed large changes among phase patterns {***θ*** ^*µ*^}. The delta-band phase dynamics *ϕ*_*i*_ (*t*) for *i* = 1, 2, …, *N* were then simulated with an increase of *D* from 0 to 1 and a step size of 0.02, and the simulated phases were labeled as 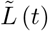 by referring to the overlap *M*_*µ*_ (*t*) [57], as follows:

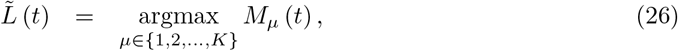

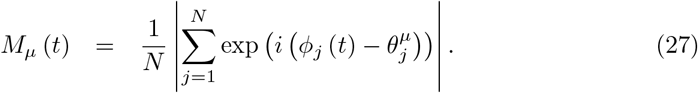

From transition 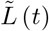, the following 3*K* statistics were extracted: the maximum 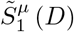 median 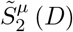, and minimum 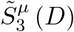 of the dwell time for *μ* = 1, 2, …, *K* with respect to each *D*, and a level 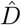 was estimated such that the root-mean-square error (RMSE) of 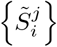 was minimized as follows:

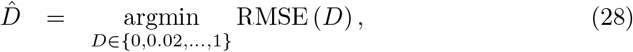

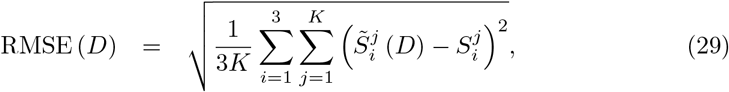

where 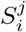 denotes the actual statistic extracted from *L*(*t*). When estimating the fluctuation level here, we set Δ*t* = 0.2 s for the sake of computational efficiency.

### Validation of the Parameter Estimation

Then, the aforementioned estimation of phase patterns {***θ***^*μ*^}, PPC connectivity ***C***^*δ*^, and the level of fluctuations *D* were validated by artificially generated phase dynamics. First, as in the previous studies [61, 62], we constructed *P* correlated patterns from a single reference pattern. From a phase pattern 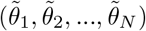 where each phase followed the uniform distribution between −*π* and *π*, we computed new pattern 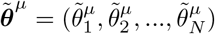as follows:

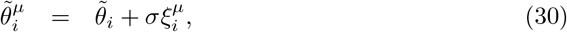

for *μ* = 1, 2, …, *P*. The variable 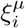 followed a normal distribution of mean 0 and standard deviation 1, and thus as *σ* approaches zero, phase patterns 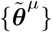 show a high correlation. The corresponding PPC connectivity 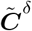 was computed using Eqs (23) and (24), and the model (Eqs (10) and (11)) combined with connectivity 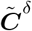 and fluctuations ***η***(*t*) was simulated. The parameter estimation was then validated in the condition of *σ* = 0.5 and *P* = 3.

### Validation of the Dynamic PPC-PAC Hypothesis

Finally, the dynamic PPC-PAC hypothesis (Fig 1) was validated by individual models (Eqs (10) and (11)) with PPC connectivity ***C***^*δ*^ and fluctuation level 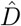, as estimated above. The delta-alpha PAC dynamics (*ϕ*_*i*_ (*t*), *r*_*i*_ (*t*)) for *i* = 1, 2, …, *N* were simulated across the subjects, and the *N* simulated CSD amplitudes (*r*_1_(*t*), *r*_2_(*t*), …, *r*_*N*_ (*t*)) were mapped to the *N* simulated EEG amplitudes *Ã*_1_(t) through observation *h* in Eq (9) as follows:

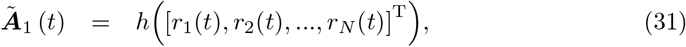

where T denotes the transposition. The simulated observations *Ã*_1_*(t)* were converted to their projection 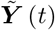 in a space with the same dimension as ***Y***(*t*). Specifically, (i) the instantaneous amplitudes *Ã*_0_(t) were computed from *Ã*1(t) around delta-band peak frequency *f*_1_, (ii) *Ã*_0_(t) were labeled as *K* states of 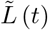, and (iii) the labeled amplitudes 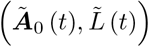 were projected to 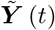 by LDA. Finally, FT surrogate data testing was performed in the same manner as for the experimental data.

## Results

We started with the condition *d* = 2, and many of the recorded brain dynamics, consisting of 63-channel scalp EEG signals ***X***(*t*), were labeled as two-dimensional metastable states with peak frequencies *f*_1_ and *f*_2_ (*n* = 101). Notably, these states were characterized by the delta- and alpha-band peak frequencies ranging from 0.1 to 4 Hz and 8 to 12 Hz, respectively (*n* = 95; Fig 3). The frequency of the fast oscillations (*f*_2_) was first estimated from the power spectrum of ***X***(*t*) (Figs 3A and 3E; S2 Fig), and was used to calculate the corresponding instantaneous amplitudes ***A***_1_(*t*) (Fig 3B). These amplitudes were converted further into corresponding instantaneous amplitudes ***A***_0_(*t*) (Fig 3C) around the frequency of the slow oscillations (*f*_1_), which was estimated from the power spectrum of ***A***_1_(*t*) (Fig 3F). Then, k-means clustering with the Calinski-Harabasz criterion [52] was applied to signals ***A***_0_(*t*), and they were labeled as *K* states of *L*(*t*) that reflected different strengths of amplitude modulation (Fig 3D). The resulting labeled signals (***A***_0_(*t*), *L*(*t*)) in Fig 3C showed significant correlations with the time courses of the modulation index [13], with the significance level of the two-sided tests being corrected for multiple comparisons using the false-discovery rate (FDR) method [63] (FDR *p* < 0.05, in the scalp sites, accounting for more than 50 electrodes; see Fig S3), and were thus characterized by different PAC strengths.

**Fig 3.**
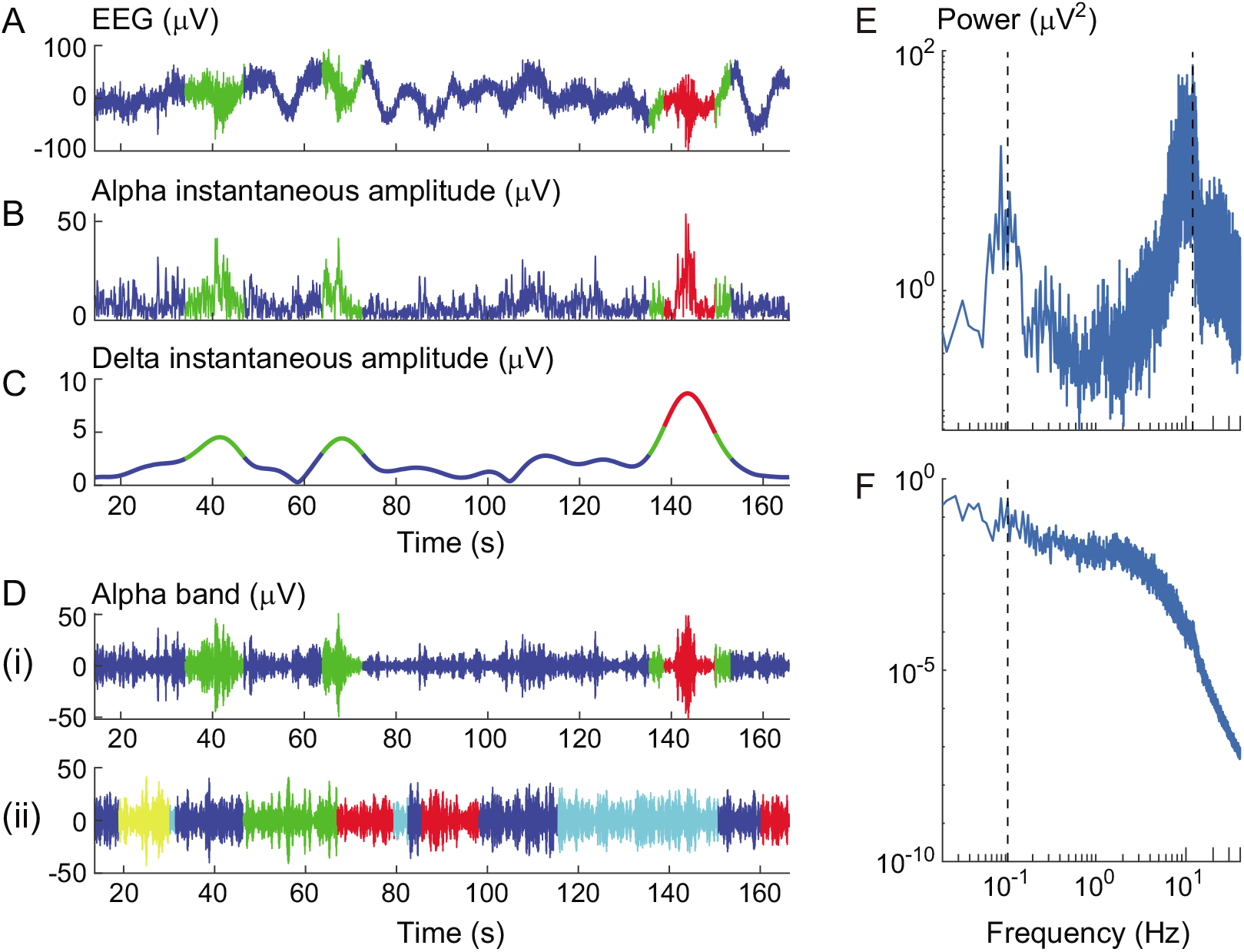
Dynamic changes in the delta-alpha PAC strength. (A) A representative raw EEG signal at the FC2 electrode. (B) The corresponding first envelope (instantaneous amplitudes) around an alpha-band peak frequency. (C) The second envelope around a delta-band peak frequency. (D) The alpha-band signals. (E) The mean power spectrum of the raw EEG signals as in panel A. (F) The mean power spectrum of the first envelopes as in panel B. The alpha-band and delta-band peak frequencies were estimated from the peaks of mean power spectra in panels E and F, respectively (see the dotted lines in panels E and F and refer to Fig S2). In panel D, (i) indicates the data corresponding to panel A, while (ii) indicates another signal of faster transition among more states obtained from an individual with a lower AQ score (the signal at electrode POz). The colors in panels A to D indicate distinct states.

To demonstrate these PAC states (two-dimensional states with PAC from *f*_1_ to *f*_2_, labeled as *L*(*t*)) as possible metastable states in the resting human brain, we projected the obtained labeled signals (***A***_0_(*t*), *L*(*t*)) to trajectory ***Y*** (*t*) in a lower-dimensional state space (Fig 4). The LDA was used to yield a space such that the projected trajectory can evolve into nearby points within each labeled state; this property is also consistent with the fixed point that can converge a trajectory into one point. Specifically, the labeled signals were projected onto a plane in the cases where the number of states was more than two (Figs 4A and 4F), and were converted into a corresponding bivariate histogram (Figs 4B and 4G); otherwise, the histogram would be a one-dimensional axis because of limitations of the LDA in this study. We calculated the maxima of the counts of bins with respect to each state, and regarded these statistics as the indices for the attracting tendencies of states. We tested whether the representative of the obtained statistics (*E*; the minimum in this study) was statistically significant, using the FT surrogate for multivariate time-series data [54] under the null hypothesis *H*_0_ where the labeled signals are linearly correlated Gaussian noise (see Materials and Methods). The surrogate data testing rejected *H*_0_ for many individual data sets under the condition of *d* = 2 (FT test, one-sided *p* <0.05, *n* = 101; Figs 4C and 4H) so that the labeled dynamics (Figs 4A and 4F; Fig 3C) were seen as a trajectory among the zero-dimensional weakly attracting states; in other words, the states of the original transition dynamics (Fig 3A) were identified as the two-dimensional metastable states. None of these data sets were rejected under condition *d* = 1, and many were not rejected for *d* = 0 (FT test, one-sided *p*>0.05, *n* = 162 for *d* = 1 and *p*>0.05, *n* = 99 for *d* = 0; see Fig S4). The majority of the other data sets were rejected for *d* = 3 (FT test, one-sided *p* < 0.05, *n* = 53). As a whole, the surrogate data testing provided experimental evidence that macroscopic brain dynamics of the resting-state large-scale network can make spontaneous transitions across two- or three-dimensional metastable states. In particular, the metastable PAC states identified here were characterized by two peak frequencies in the delta and alpha bands (*n* = 95). We represented each of these delta-alpha PAC states by a vector composed of mean values of the labeled signals (i.e. the delta-band instantaneous amplitudes depicted in Fig 3C) over time, namely, by a 63-dimensional vector of mean PAC strengths (Figs 4D and 4I; Fig S3).

**Fig 4.**
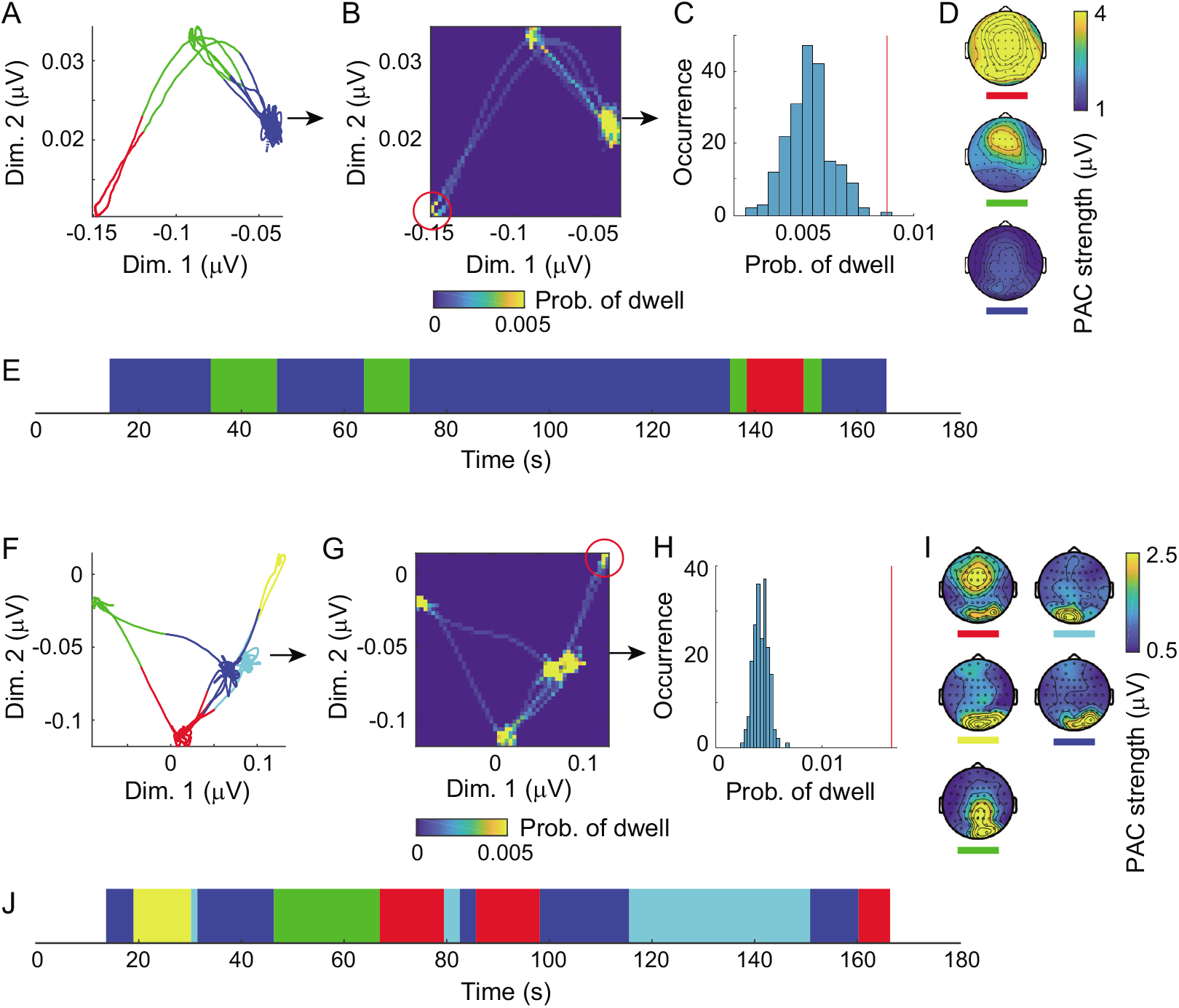
Transition dynamics among delta-alpha PAC states in a lower-dimensional space. (A to E) Representative delta-alpha PAC dynamics for an individual with higher attention-related AQ subscores (refer to Figs 3A to 3D(i)). (F to J) Representative dynamics for an individual with lower scores (refer to Fig 3D(ii)). (A, F) The trajectory of labeled signals in a plane. (B, G) The corresponding bivariate histograms. (C, H) Surrogate data testing under a condition of *d* = 2. (D, I) The resulting delta-alpha PAC states (represented by the mean PAC strengths). (E, J) Transitions between the identified PAC states. Surrogate data testing was applied to the density of points indicated by the red circles in panels B and G and the red lines in panels C and H, and the null hypothesis *H*_0_ in condition *d* = 2 was rejected (C and H). The delta-alpha PAC dynamics tended to stay in a state for a longer time and to visit a lower number of states in individuals with higher AQ subscores for attention to detail and attention switching (compare E with J). The colors in panels A, E, F, and J indicate distinct states, as depicted in D and I.

Delta-alpha PAC states (Figs 4D and 4I) were categorized into four groups across individuals (*n* = 95; Fig 5). First, we converted these states into modified Z-scores [64] to standardize them robustly among individuals, with each data set being subtracted by the median and divided by the median absolute deviation instead of the mean and the SD, respectively, and all the data were subsequently multiplied by 0.6745 [64]. The obtained Z-scores were concatenated across states and individuals, and the resulting dataset was regarded as the data in a 63-dimensional feature space. In this space, we conducted principal component analysis (PCA) and applied a permutation test to 63 PCs by shuffling the dataset 200 times across the channel with respect to each component. The first four PCs significantly explained variance (one-sided, Bonferroni-corrected *p* < 1.58×10^−4^, total explained variance 81.6 %; Fig 5A). Eigenvectors of these four PCs were then mapped as the topographies and categorized according to the regional distribution of the amplitude modulation in the occipital lobe, parietal lobe, and lateral and bilateral distributions in the occipital lobe (Fig 5B).

**Fig 5.**
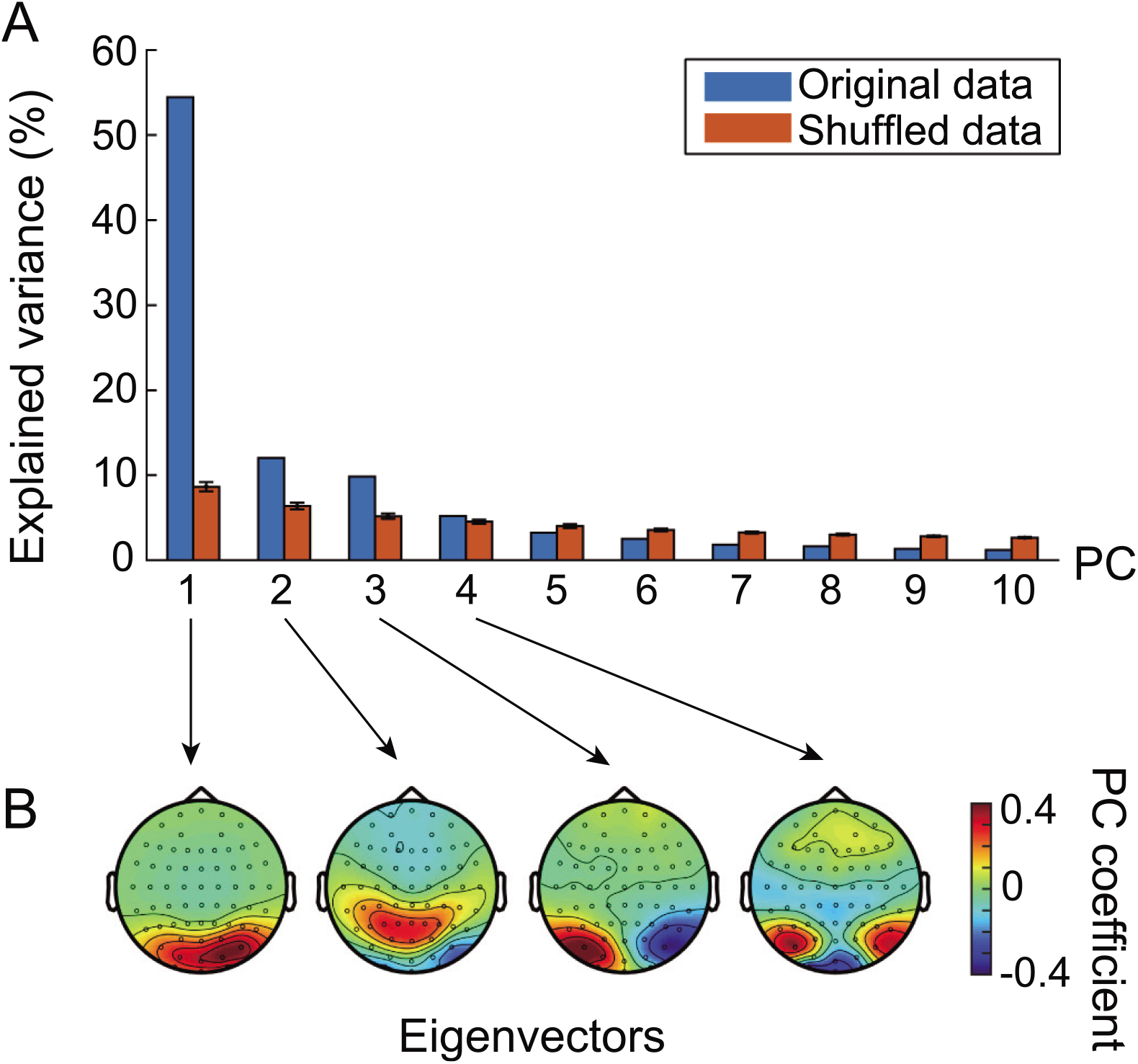
The four groups of consistent delta-alpha PAC states across individuals. (A) PCs of across-individual states. (B) Eigenvectors of the first four PCs. The variance explained by the first four PCs was significant, and accounted for 81.6 % of total variance. The dataset used here was a set of the modified Z-scores of mean PAC strengths that were concatenated across states and individuals.

The dynamics of transitions between the delta-alpha PAC states (Figs 4E and 4J), as identified in this study, showed correlations with the two AQ subscores of ‘attention to detail’ and ‘attention switching’ (*n* = 52; Fig 6). From the transition dynamics, we calculated the intervals between transitions (for which uncertain intervals at both edges were excluded) and obtained the following candidate statistics: the maximum, median, and minimum of the dwell time. These statistics, in addition to the number of states and the alpha- and delta-band peak frequencies estimated above, were regarded as test statistics (*x*) and were paired with the following five AQ subscores (*y*): social skills, attention to detail, attention switching, communication, and imagination. For these 30 pairwise statistics, we used multiple comparison tests with Pearson’s correlation coefficients. The maximal dwell time showed a significant positive correlation with the attention-to-detail score (*r* = 0.456, two-sided, Bonferroni-corrected *p* < 0.0013; Fig 6A, the effects of remaining variables in *x* on *y* were partially adjusted). Next, we conducted a post-hoc test of the multiple correlation coefficient using a linear regression model in which the attention-switching score was regarded as a dependent variable and was regressed against two statistics: the number of states and the alpha-band peak frequency (Fig 6B), with these being selected because of weak significant correlations with the attention-switching score (*r* = − 0.283, two-sided, uncorrected *p <* 0.06 for the number of states; and *r* = 0.321, two-sided, uncorrected *p* < 0.03 for the alpha-band peak frequency; Figs 6C and 6D, the effects of the remaining variables in *x* on *y* were partially adjusted). The resulting linear combination showed significant correlation with the attention-switching score (*F* (2, 49) = 4.91, *r* = 0.409, *p <* 0.02), and factor loadings of this linear sum on the number of states and the alpha-band peak frequency (i.e. the correlation coefficients) were − 0.614 and 0.666, respectively (Fig 6B). The results indicated that in individuals with the ability to maintain a stronger focus on attention to detail and less attention switching, the delta-alpha PAC dynamics tended to stay in a particular state for a longer time, to visit a lower number of states, and to oscillate at a higher alpha-band peak frequency, thereby providing evidence on how autistic-like traits may be associated with the metastable human brain.

**Fig 6.**
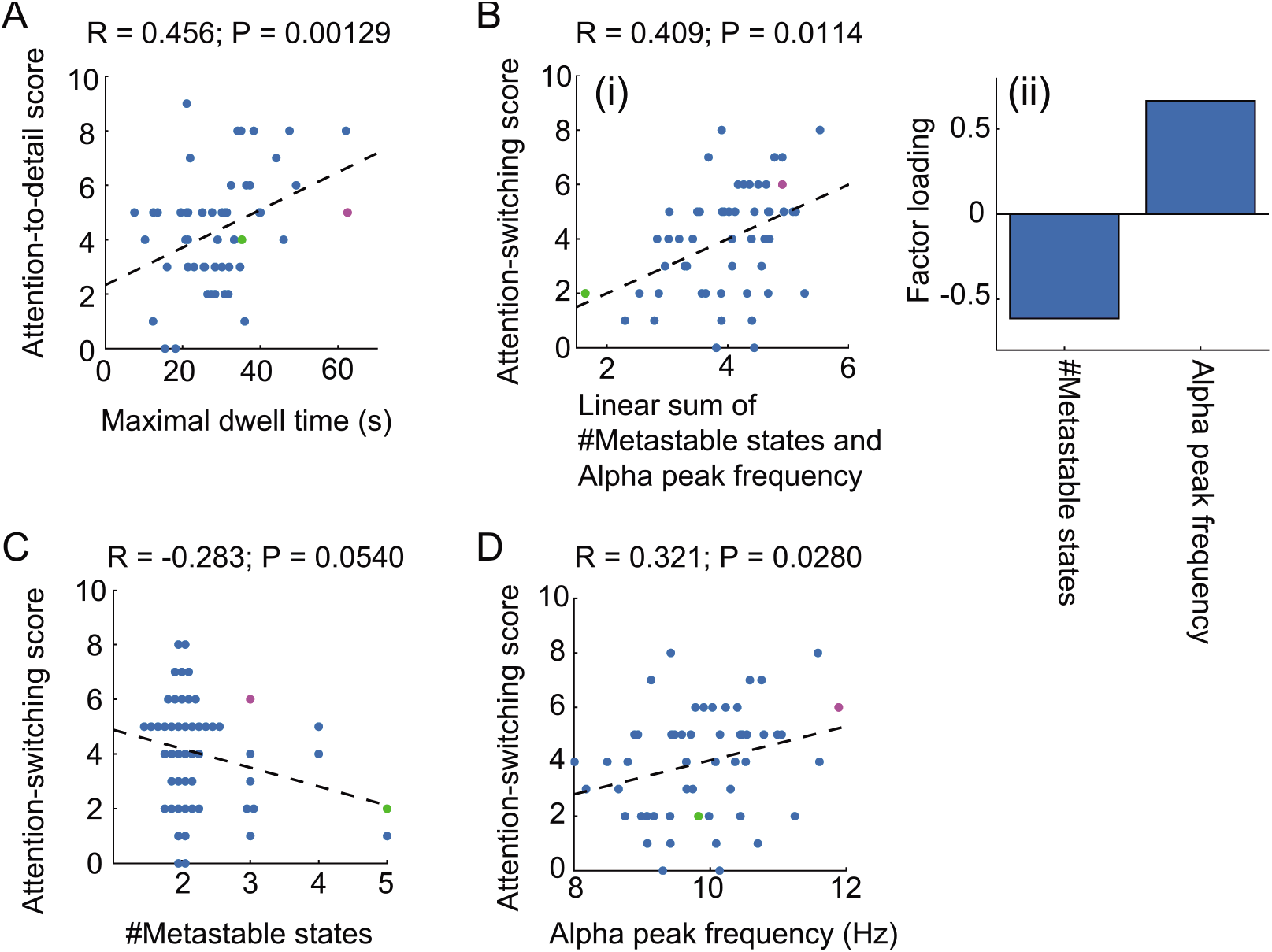
Correlations between delta-alpha PAC dynamics and attention-related AQ subscores. (A) Scatter plot of the attention-to-detail score against maximal dwell time. (B) Scatter plot of the attention-switching score against the linear sum of the number of states and the alpha-band peak frequency (i) with corresponding factor loadings (ii). (C, D) Scatter plots of the attention-switching score against the number of states and the alpha-band peak frequency, respectively. In each panel, the circles in magenta and green correspond to the representative individual delta-alpha PAC dynamics, as depicted in Figs 4A to 4E and Figs 4F to 4J, respectively. The dotted line in each panel indicates the fitted regression line.

We modeled the metastable delta-alpha PAC dynamics (*n* = 95) to validate the dynamic PPC-PAC hypothesis (Materials and Methods; Fig 1). The model consisted of delta-alpha PAC oscillators {*ϕ*_*i*_(*t*), *r*_*i*_(*t*)}, PPC-PAC connectivity (***C***^*δ*^, ***C***^*αδ*^), and fluctuations *D****η***(*t*). More specifically, we made connections among the delta-band phases (***C***^*δ*^: PPC connectivity), from delta-band phases to alpha-band amplitudes (***C***^*αδ*^: PAC connectivity) and from fluctuations to delta-band phases (*D*: fluctuation level), such that synchronization, amplitude modulation, and state transition could occur via the PPC and PAC. The PPC connectivity (***C***^*δ*^) and the level of fluctuations (*D*) were estimated from the data for each individual (Fig 7); this parameter estimation was validated by the artificially generated phase dynamics (Fig S5). On the other hand, the PAC connectivity (***C***^*αδ*^) was set to arbitrary values, because our hypothesis was that state transition occurs according to dynamic changes in PPC strengths (Fig 1). Note that a part of the present model was composed of only delta-band phases (Eq (11)), with the PPC being equivalent to the Kuramoto model subjected to noise [23]. In this study, the metastable PAC dynamics were modeled as the fluctuation-induced transitions between multistable attractors (for the scenario of multistability, see [58]).

**Fig 7.**
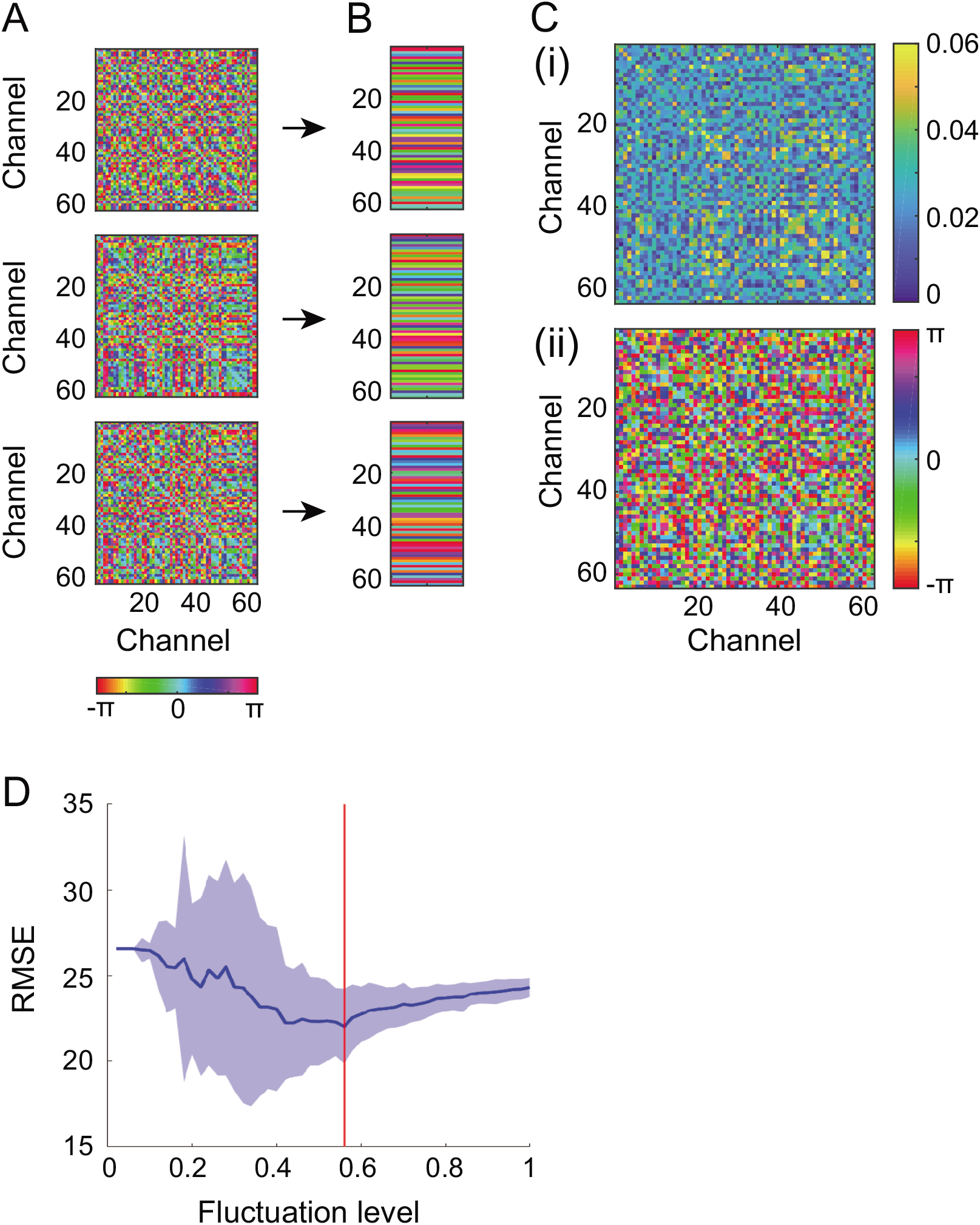
Estimation of PPC connectivity and the level of fluctuations from delta-band phase dynamics. (A) Mean phase differences 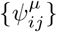 between every pair of delta-band CSD phases with respect to each delta-alpha PAC state *µ*. (B) The corresponding phase patterns {***θ***^*µ*^} (regarded as the synchronous states in the model) (C) The estimated PPC connectivity ***C***^*δ*^ as a complex-valued matrix with absolute part 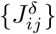 (i) and argument part 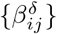 (ii). (D) The estimated fluctuation level 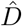 The synchronous states (B) combined with the Kuramoto model resulted in PPC connectivity (C), and the Kuramoto model with PPC connectivity was used for estimation of the fluctuation level (D). The data used in this Figure correspond to Figs 3A to 3D(i) and Figs 4A to 4E.

First, we computed the CSD [42, 55] from raw EEG signals to reduce the volume-conduction effect on the estimation of instantaneous phases, and then estimated the phase patterns {***θ*** ^*µ*^} from the 63-channel CSD signals ***I***_m_(*t*) (Figs 7A and 7B). Specifically, we extracted the instantaneous phases ***θ***(*t*) from ***I***_m_(*t*) around the delta-band peak frequency *f*_1_, and labeled ***θ***(*t*) as transition *L*(*t*) (refer to Figs 4E and 4J). We then computed the phase differences 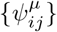 from labeled phases (***θ***(*t*), *L*(*t*)) with respect to each state *µ* (Fig 7A), and estimated the phase patterns {***θ***^*µ*^}, which were regarded as the synchronous states in the model.

Next, we estimated the PPC connectivity ***C***^*δ*^ and the level of fluctuations *D* (Figs 7C and 7D). Synchronous states {***θ***^*µ*^} obtained above were applied to the Kuramoto model (Eq (11)) composed of delta-band phases {*ϕ*_*i*_(*t*)}, and they were converted into PPC connectivity ***C***^*δ*^ (Fig 7C) [57]. We gradually increased the level (*D*) of fluctuations to phases, simulated the corresponding models, and generated the transitions for each level. The resulting set of transitions was quantified by the maximum, median, and minimum of the dwell time with respect to each fluctuation level, and from these statistics and those obtained from the data, we calculated the RMSE. We repeated this calculation 100 times and chose the fluctuation level 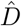 minimizing the RMSE (Fig 7D).

Then, we simulated the individual delta-alpha PAC dynamics (*n* = 95) as modeled above (Fig 8). By calculating the overlaps *M*_*µ*_(*t*) (Eq (27)) every time step and with respect to each state *µ* (Figs 8A and 8E), we observed from the model that the phase differences *{ϕ*_*i*_(*t*) − *ϕ*_*j*_(*t*)} changed dynamically among synchronous states {***θ***^*µ*^}, and that the alpha-band amplitudes {*r*_*i*_(*t*)} were oscillatory at the delta-band frequency *f*_1_. These oscillatory amplitudes were applied to observation *h* in Eq (31), so that the simulated EEG amplitudes ***Ã***_1_(*t*) were obtained. We then computed corresponding instantaneous amplitudes ***Ã***_0_(*t*) around *f*_1_, labeled ***Ã***_0_(*t*) as transition 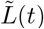 by referring to the overlaps (Eqs (26) and (27)), and projected the labeled signals 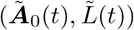 to trajectory 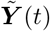 in a lower-dimensional space (Figs 8B and 8F). This space was converted into a corresponding histogram (Figs 8C and 8G), which we used to conduct surrogate data testing in the same manner as for the experimental data (***A***_0_(*t*), *L*(*t*)). For simulated delta-alpha PAC dynamics, the surrogate data testing rejected *H*_0_ under the more in *d* = 2 compared to *d* = 1 (FT test, one-sided *p <* 0.05, *n* = 74 out of 95 for *d* = 2 and *p<*0.05, *n* = 45 out of 95 for *d* = 1; Figs 8D and 8H, Fig S6). A hypothesis test for the difference in two population portions indicated that the portion of the rejected participants was significantly higher in *d* = 2 than in *d* = 1 (*p <* 0.001) Overall, these simulation results (Fig 8) were consistent with the empirical counterpart (Fig 4), providing evidence for the dynamic PPC-PAC hypothesis.

**Fig 8.**
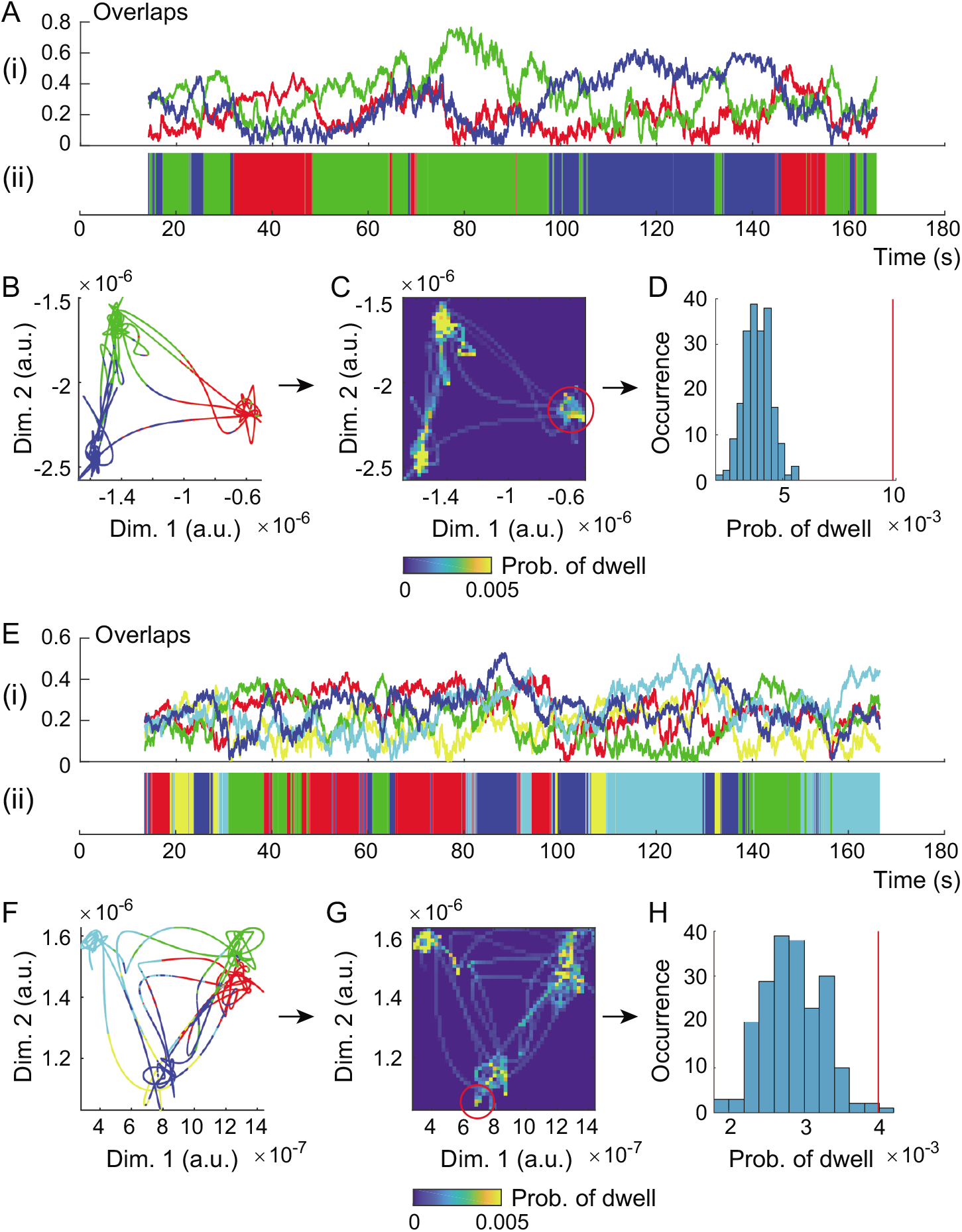
Simulation of delta-alpha PAC dynamics by a coupled oscillator system driven by spontaneous fluctuations. (A to D) The representative simulated delta-alpha PAC dynamics for an individual with higher attention-related AQ subscores (refer to Figs 4A to 4E). (E to H) Representative dynamics for an individual with lower scores (refer to Figs 4F to 4J). (A, E) Time courses of overlaps (i) and the corresponding labels (ii) among delta-alpha PAC states. (B, F) The trajectory of labeled signals in a plane. (C, G) The corresponding bivariate histograms. (D, H) Surrogate data testing under condition *d* = 2. Surrogate data testing was applied to the density of points indicated by the red circles in panels C and G and the red lines in panels D and H, and the null hypothesis *H*_0_ in condition *d* = 2 was rejected (D and H). The model showed consistent results with the data analysis, evidence of the dynamic PPC-PAC hypothesis.

Finally, we attempted to predict the delta-alpha PAC dynamics with a temporally decreasing fluctuation level *D*(*t*) (Fig 9). By calculating the overlaps every time step in this simulation, we observed that one of the delta-alpha PAC states was stabilized, so that the transition dynamics qualitatively changed into the dynamics in a single state (Fig 9A) as the fluctuation level decreased (Fig 9B). This appearance depended on the initial condition of the system. Moreover, such a qualitative change from multiple states to one state was viewed as a shrinking of the trajectory in the state space (Fig 9C). We generated the trajectories of the system under different initial conditions in a space composed of the overlaps. The trajectories were projected onto planes, from which we observed that the spaces filled by the transition dynamics can include the three states as their subsets (Fig 9C).

**Fig 9.**
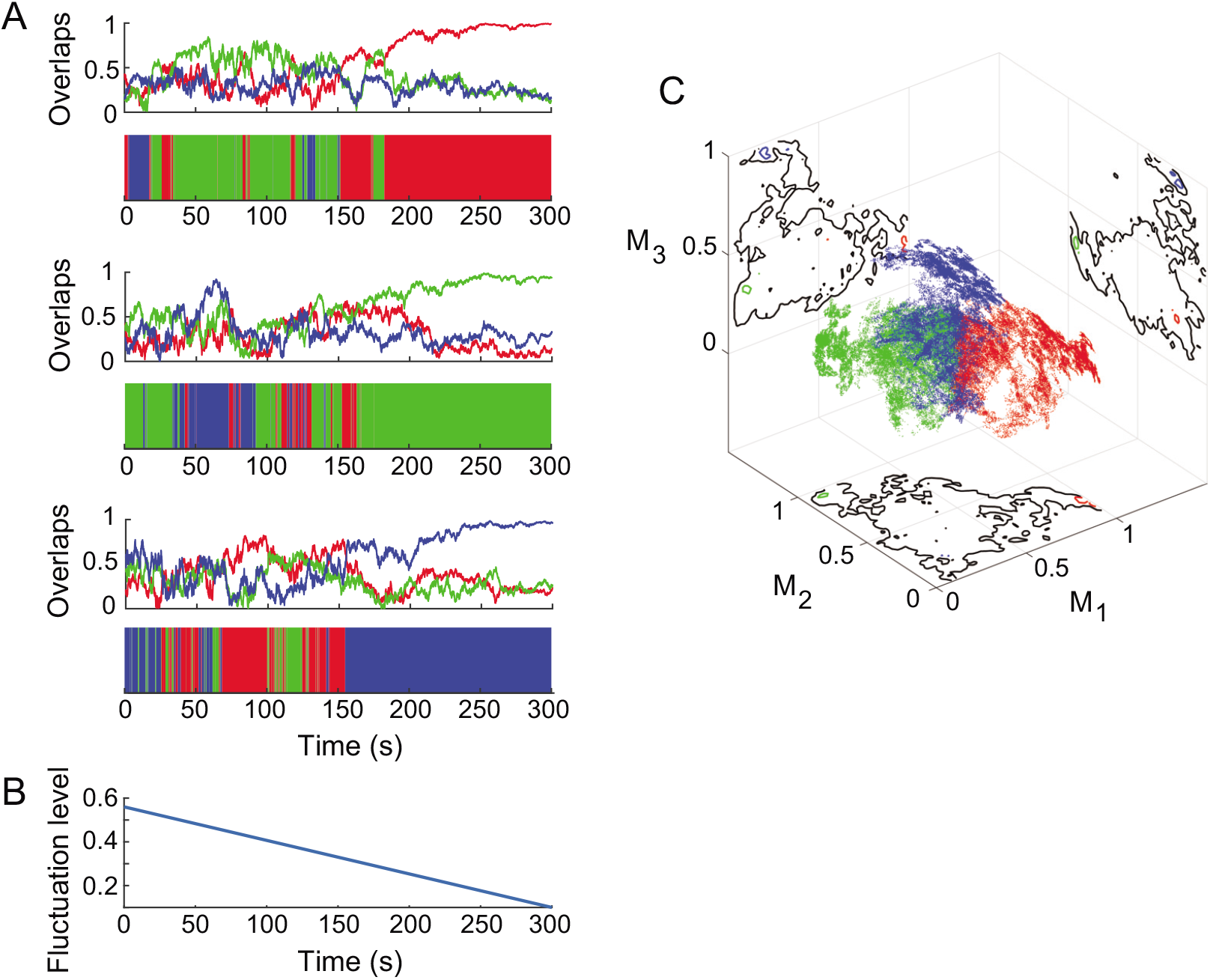
Shrinking of the simulated delta-alpha PAC dynamics with a temporally decreasing fluctuation level in the phase space: the qualitative change from the transition dynamics to the dynamics in a single state. (A) Time courses of the overlaps with their labeled sequences under different initial conditions in cases where the dynamics can converge into one of three states. (B) The time course of the fluctuation level. (C) The trajectories of overlaps in the phase space with their projections. The contour plots on projections in panel C indicate that the spaces filled by transition dynamics (black lines) can include the three states (red, green, and blue lines) as their subsets. The data used in this Figure correspond to the individual with higher attention-related AQ subscores depicted in Figures 3A to 3D(i), 4A to 4E, 7 and 8A to 8D.

## Discussion

In this study, we developed a data-driven approach to label observed metastable dynamics as the underlying *d*-dimensional states. The method was applied to channel scalp EEG signals recorded from 130 healthy humans in an eyes-closed resting condition (*n* = 162 in total). The observed signals were labeled as metastable states with a dimension larger than one, such that PAC could occur hierarchically, in particular with a dimension of two, corresponding with the delta- and alpha-band peak frequencies (*n* = 95; Fig 3). Then, the dynamics of the transitions between the delta-alpha PAC states (Fig 4), which were categorized into four groups across individuals (Fig 5), showed correlations with the AQ subscales of attention to detail and attention switching (Fig 6). Finally, we qualitatively reproduced the obtained results in a coupled oscillator system driven by spontaneous fluctuations (Figs 7 and 8) with prediction (Fig 9), to validate the hypothesis that the dynamic changes in PAC strengths can be attributed to changes in the strengths of PPC, that is, to dynamic functional connectivity in an electrophysiological sense (Fig 1).

Many studies show that neural activity exhibits oscillations whose amplitudes change rhythmically over time [6, 12, 13, 65, 66]. A possible mechanism for this amplitude modulation is PAC, in which the phases of slow oscillations interact with the amplitudes of faster oscillations such that local and global computations in the large-scale network can cooperate [12, 13]; the PAC takes various forms depending on events such as visual and auditory tasks [65, 66]. In the present work, we identified a possible link of the PAC for resting-state EEG dynamics to the two-dimensional metastable states, which were also characterized by slow and fast timescales (Figs 1 to 4). Some modeling studies have shown evidence for a torus-related PAC [18, 19], such as the study of Sase et al., which analyzed a model composed of excitatory and inhibitory networks with dynamic synapses and revealed that amplitude-modulated dynamics can emerge from a trajectory into the torus or the closed curve, with these being mediated by bifurcation [18]. Hence, it is suggested that attractors in the resting brain play a functional role in generating cooperative dynamics over the large-scale network and could be a torus, in which spontaneous events are effectively processed by the utilization of multiple neural timescales.

A two- or three-dimensional state was identified as a possible metastable state underlying the resting-state EEG signals of most individuals (refer to Figs 1 to 4). This result implies that macroscopic dynamics in the human brain can follow the oscillatory hierarchy hypothesis stating that slower and faster oscillations interact hierarchically via the PAC [35, 67]. Lakatos et al. showed experimental evidence of EEG hierarchical organization: delta-band phases modulated amplitudes of fast oscillations, which further made a spontaneous PAC connection to another faster oscillatory component [35]. We suggest that the resting human brain could utilize attractors with multiple timescales, so that a variety of events are spontaneously organized in a hierarchy of macroscopic neural oscillations.

Not only were the amplitudes of resting-state EEG signals rhythmic, but the strengths of the PAC (as obtained after two-time computations of instantaneous amplitudes) exhibited dynamic changes such that transitions between two-dimensional states could occur (Fig 4). Previous studies showed that the PAC strength changed dynamically over time, and transiently in response to sensory and cognitive events [12, 68–71]. Such dynamic coupling was observed during cognitive behavior in a T-maze task [68], learning [69], attentional allocation [70], and motor preparation [71]. Using a spatial-cuing task, Szczepanski et al. found PAC modulation-dependent attentional behavior in which the modulation strength was negatively correlated with the reaction time on a trial-by-trial basis [70]. Moreover, Kajihara et al. showed evidence that delta-alpha PAC dynamically occurs to mediate the global-to-local computation in motor preparation, such that the delta-band synchrony makes a direct link with the alpha-band amplitudes via the PAC [71]. These results may be supported by the conventional view of the task-dependent PAC [65, 66]: Voytek et al. reported that fast oscillations were strongly coupled with a slow oscillatory component via PAC during a visual task, and that this coupling weakened during an auditory task so that PAC with another slow oscillatory component could occur [66]. In recent years, it has been suggested that dynamic PAC plays a role in modulating the dynamics of the large-scale network, doing so more effectively than coupling with static modulation [12].

To the best of our knowledge, our finding of dynamic PAC, as realized by the transition between the metastable states, is the first experimental report of this phenomenon. Crucially, we identified the transitions between delta-alpha PAC states (Fig 4), which were further categorized into four groups across individuals (*n* = 95; Fig 5). Previous studies showed evidence from resting-state functional magnetic resonance imaging (fMRI) signals that large-scale subnetworks with different functional connectivity, termed ‘resting-state networks’, are consistent across individuals [72, 73], and that these consist of the following components: the default model network, the executive control network, the salience network, the dorsal attention network, and networks related to auditory, sensorimotor, and visual functions [73]. In recent years, it has been suggested that such networks are linked to the underlying electrophysiological oscillations [74, 75]. With respect to each network, Mantini et al. showed correlations between slow fluctuations in the blood-oxygen-level-dependent (BOLD) signal and EEG power variations of different brain rhythms, including delta and alpha rhythms [74]. Moreover, Britz et al. identified four resting-state networks from BOLD signals combined with the transition dynamics of EEG scalp potentials [75], referred to as EEG microstates [24–27], and a previous study likewise showed four network modules that were highly consistent across subjects [72]. These results inspired attempts to detect the large-scale functional network using only EEG data [76]. Moreover, the regional specificity of PAC has also been reported [66, 70], as well as the lateralization of PAC strengths [70]. Thus, macroscopic neural oscillations with multiple timescales in the resting human brain, identified as the delta-alpha PAC states in this study, could be the electrophysiological signatures of resting-state networks.

Our main finding is the AQ-related behavioral correlates of delta-alpha PAC dynamics, namely, the correlation with the two AQ subscales of attention to detail and attention switching (Fig 6). In fact, slower neural oscillations are suggested to be dynamically entrained by rhythmic input from external sensory events [12, 14, 77]. Lakatos et al. showed that delta-band oscillations selectively entrained to the rhythm of attended visual and auditory stimuli, thereby providing evidence of the neural entrainment to attention by which the brain encodes task-relevant events into preferred delta-band phases [14]. On the other hand, alpha-band oscillations have been suggested to play an inhibitory role by effectively gating top-down processing [78]. Previous studies showed that the alpha-band power decreased in the hemisphere contralateral to attended visual stimuli, whereas it exhibited an increase in the ipsilateral hemisphere (refer to Fig 5) [78]; this is evidence for attention-induced alpha-band lateralization that gates the flow of top-down information into task-irrelevant regions [78, 79]. Alpha-band activity are dominantly observed in the resting brain, in particular in the occipital region [7], and the alpha-band peak frequency depends on age and cognitive performance [7], which shows inter-individual variability [80]. Moreover, a recent study reported atypical neural timescales for individuals with autism spectrum disorder (ASD) [81], on the basis of the fact that the heterogeneity of timescales in the brain could be a basis for functional hierarchy [82]. Watanabe et al. found shorter neural timescales in sensory/visual regions and a longer timescale in the right caudate for individuals with a higher severity level of ASD [81]. Together, it is suggested that attractors in the resting human brain generate individual delta-alpha PAC dynamics that selectively encode spontaneous events by utilizing attention. Individual macroscopic dynamics in the brain, as identified here, and which tend to stay in a state for a longer time, to visit a lower number of states, and to oscillate at a higher alpha-band frequency in individuals with a stronger preference for specific events (Fig 6), might be a neural signature of the autism spectrum, covering both typical and atypical development.

Although we filtered out lower frequency components below 0.1 Hz offline, the delta-band oscillations (0.1–4 Hz) reported here may still be affected by infra-slow fluctuations (0.01–0.1 Hz) during rest. Such spontaneous fluctuations have been observed by various neuroimaging methods such as EEG and fMRI [83, 84], and a previous study reported that spontaneous infra-slow scalp potential fluctuations are correlated with BOLD fluctuations of resting-state networks [85]. In this study, the lower bound of the delta-band frequency was set to the lower bound of the spectral frequency of the first envelopes, i.e., 0.1 Hz. Dependence of the results on this parameter should be investigated in future work.

Recently, atypical transition dynamics of the resting large-scale network were identified as ASD symptoms [86]. By applying energy-landscape analysis [87] to the fMRI signals of resting-state networks, Watanabe and Rees showed that neurotypical brain activity transited between two major states via an intermediate state, and that the number of these transitions was lower due to the unstabilization of the intermediate state for the individuals with a higher severity level of ASD [86]. Such dynamics-behavior associations were linked to functional segregation. In this study, we generated the energy-like landscape of resting-state EEG dynamics by utilizing multi-step envelope analysis (Fig 2), so that the underlying oscillatory attractors could transform into the fixed points (Fig 4), and found a similar dynamics-behavior association between the dwell time of delta-alpha PAC state transitions and the attention-to-detail AQ subscale (Fig 6A). Hence, the individual delta-alpha PAC dynamics could be the electrophysiological signature of an atypical balance in functional organization.

What kind of mechanisms can underlie the dynamic PAC and enable transitions between attractors? One possible mechanism is the metastability (or called criticality in a similar sense) that is suggested to play a role in maintaining a dynamic balance of integration and segregation of brain functions across multiple spatiotemporal scales [4, 34, 88]. Such dynamic organization was fruitfully discussed from viewpoints of both models and experimental data by Tognoli and Kelso [34]. By introducing an extended Haken-Kelso-Bunz model [88] and actual neurophysiological and behavioral data [34], they illustrated that phase dynamics in the brain can utilize both tendencies of dwells to be in synchrony and escapes into non-synchronous patterns, and associated this fact with the abilities of the brain (integration and segregation) in the theory of coordination dynamics [34, 88]. Similar dynamics were previously observed in the resting-state neural signals of EEG [26, 28, 29], fMRI [89], and functional multineuron calcium imaging [90] aimed at generating a better mathematical model of individual brains [1–5, 18, 20, 21, 23, 84, 91]; there is ongoing debate whether spontaneous neural activity originates from a deterministic dynamical system [88] that may yield chaotic itinerancy [92], or our present standpoint, a random dynamical system driven by spontaneous stochastic fluctuations [93–95] (see [58] for possible scenarios). Together, it is suggested that individual delta-alpha PAC dynamics at rest (which could relate to previous studies reporting that delta-alpha PAC occurred in preparation for a task [71] and during decision making [65]) utilize metastability to organize spontaneous events in a hierarchy of macroscopic oscillations with multiple timescales.

Here, on the basis of our dynamic PPC-PAC hypothesis validated by the empirical and modeling counterparts (Fig 1; Figs 4 and 8), we posit that the dynamic changes in delta-alpha PAC modulation are attributed to the dynamic functional connectivity in an electrophysiological sense. Dynamic functional connectivity is referred to as the functional connectivity of the large-scale network with dynamically changing temporal correlation [96], and has been regularly observed in resting-state fMRI signals with behavioral and cognitive relevance [89, 97]. In a similar sense, resting-state EEG experiments showed that the dynamics of the large-scale network transited among a repertoire of synchronized states [8, 28, 29]. We applied this view to the large-scale network of slow oscillations, taking into account its neuromodulatory influences on a faster oscillatory component, and validated the resulting dynamic PPC-PAC hypothesis using an extended version of the Kuramoto model (see Figs 1, 7, and 8). By analyzing a model of local networks with heterogeneity near the onset of synchrony, a relevant modeling study demonstrated that transient synchrony of the large-scale network can organize the routing of information flow [98]. In the present study, we detected the synchronous states by referring to the transitions of delta-alpha PAC dynamics (Figs 4E and 4J) to support the dynamic PPC-PAC hypothesis. However, the changes in PPC connectivity would be better characterized by drifts of the delta frequency that might occur from transitions. Dynamic and transient delta-alpha PAC, as identified in this study, may originate from the coupling between delta-band phases, and the relationship of this transient PAC with transient synchrony remains to be elucidated.

We should remark on the observation operator *h* in Eqs (9) and (31) that connected the simulated CSD dynamics to the experimental EEG data. The method (Fig 2) was developed such that the estimation of inter-regional properties between signals was not necessarily required, and accordingly it was directly applied to the scalp EEG signals. While it might seem natural to apply the CSD to the empirical counterpart as well, we wanted to avoid generating possible artifacts due to the CSD estimation [56]. However, amplitude modulation of apparent frequencies could affect the results when multiple uncoupled sources coexist in the brain. Limitations of the metastable states clustering developed here should be clarified in future work.

Finally, we observed shrinking of the transient PAC dynamics with a temporally decreasing fluctuation level from the model (Fig 9). This result could relate to the reduction in trial-to-trial variability of cortical activity that can occur after the stimulus onset, such that the spontaneous and task-evoked brain activity interplay in a complex manner [99]. Such a phenomenon was previously observed from the spikes of single neurons [100], and was recently demonstrated by a model including local and global cortical networks at multiple spatiotemporal scales [101]. Thus, the present model combined with the resting-state EEG data could have the potential for predicting task-relevant events; for example, identifying a parameter that can facilitate a dynamic balance in the typical and atypical neural activity of the large-scale network, which might be helpful for mitigating the severity level of ASD, so that faster transition dynamics among more states can appear during rest.

Taken together, we reported the first experimental evidence that (i) metastable states in the resting human brain can be two- or three-dimensional; (ii) that their dynamics can be metastable delta-alpha PAC dynamics; and (iii) their functional role is associated with autistic-like traits. These empirical findings were then combined with the modeling counterpart, to provide evidence that (iv) such dynamic and functional PAC may originate from dynamic functional connectivity. We suggest that the metastable human brain organizes spontaneous events dynamically and selectively in a hierarchy of macroscopic oscillations that interact in a cooperative manner, and that such dynamic organization might encode a spectrum of individual traits covering both typical and atypical development. Our findings on the metastable human brain and its association with autistic-like traits may be further corroborated by the following research: (i) the brain of ASD subjects during rest [86, 102, 103] to verify our present findings from healthy subjects; (ii) the brain during transcranial magnetic stimulation [32, 104] and closed-loop control by neurofeedback [79] to manipulate individual traits; and (iii) the brain during a task to understand the relationship between spontaneous and task-evoked dynamics from the viewpoint of the attractors that might underlie the human brain [99].

## Acknowledgments

KK was supported by a research grant from TOYOTA Motor Corporation. We are grateful to Ms. Yoko Noguchi for help with acquisition of EEG signals and AQ scores, and to Dr. Yuka O. Okazaki for helpful discussion.

## Author Contributions

TS contributed to conceptualization, data curation, formal analysis, investigation, methodology, software, validation, visualization, writing - original draft, and writing - review & editing. KK contributed to conceptualization, data curation, formal analysis, funding acquisition, investigation, methodology, project administration, resources, supervision, validation, and writing - review & editing.

## Data Availability

All relevant data and code are available in OSF. https://osf.io/29qb5/doi10.17605/OSF.IO/29QB5

