## Supplemental figure S1-6 for "The metastable brain associated with autistic-like traits of typically developing individuals"

### Supporting Information

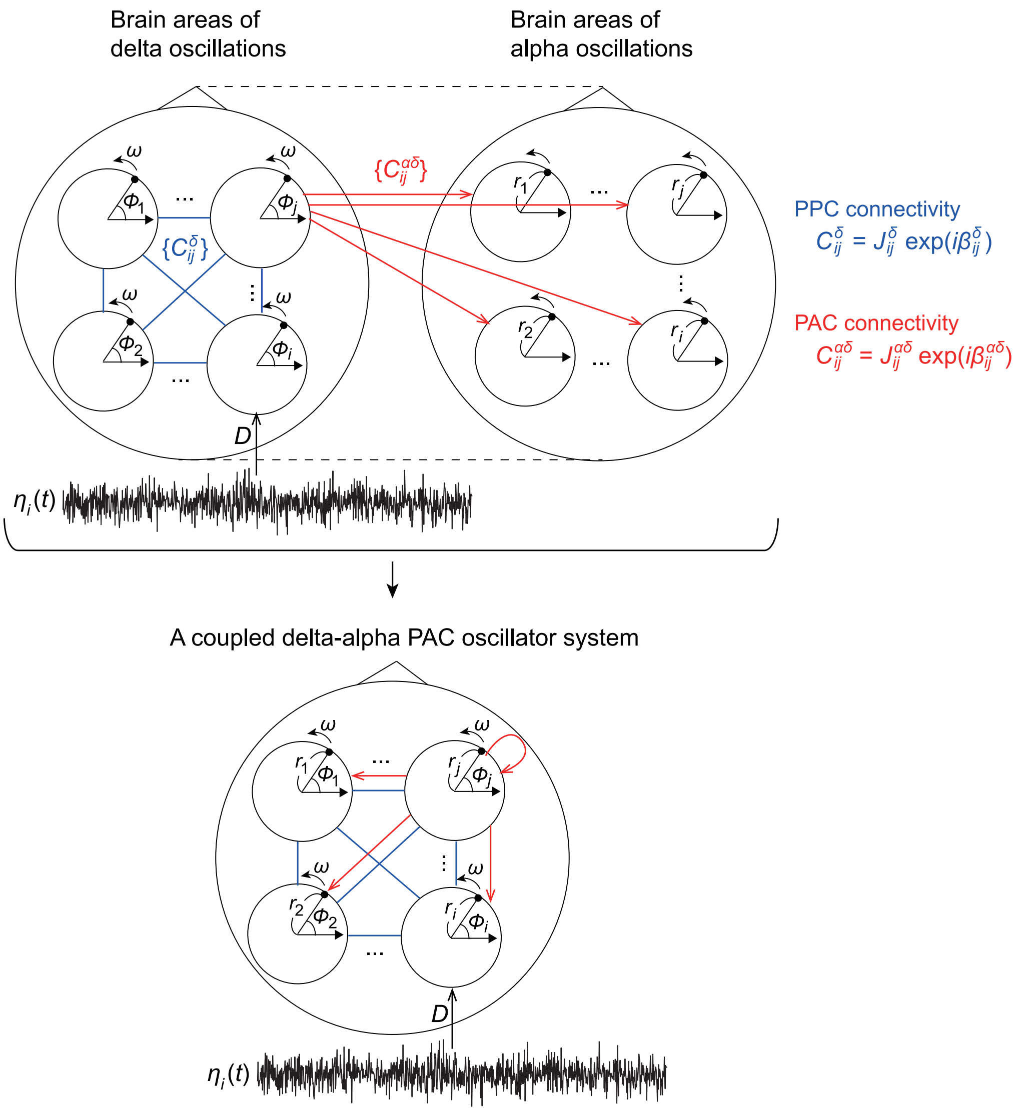


**Fig S1. The model of a coupled oscillator system of delta-alpha PAC driven by fluctuations.** The model comprised $N$ PAC oscillators whose phases $\{\phi_i(t)\}$ and amplitudes $\{r_i(t)\}$ corresponded to delta- and alpha-band activity, respectively. The phase $\phi_j(t)$ interacted with $\phi_i(t)$ and $r_i(t)$ via the PPC connectivity $C^{\delta}_{ij} = J^{\delta}_{ij} \exp(i\beta^{\delta}_{ij})$ and the PAC connectivity $C^{\alpha\delta}_{ij} = J^{\alpha\delta}_{ij} \exp(i\beta^{\alpha\delta}_{ij})$. The phase $\phi_i(t)$ was driven by fluctuation $\eta_i(t)$ with the level $D$ for $i = 1, 2, ..., N$.

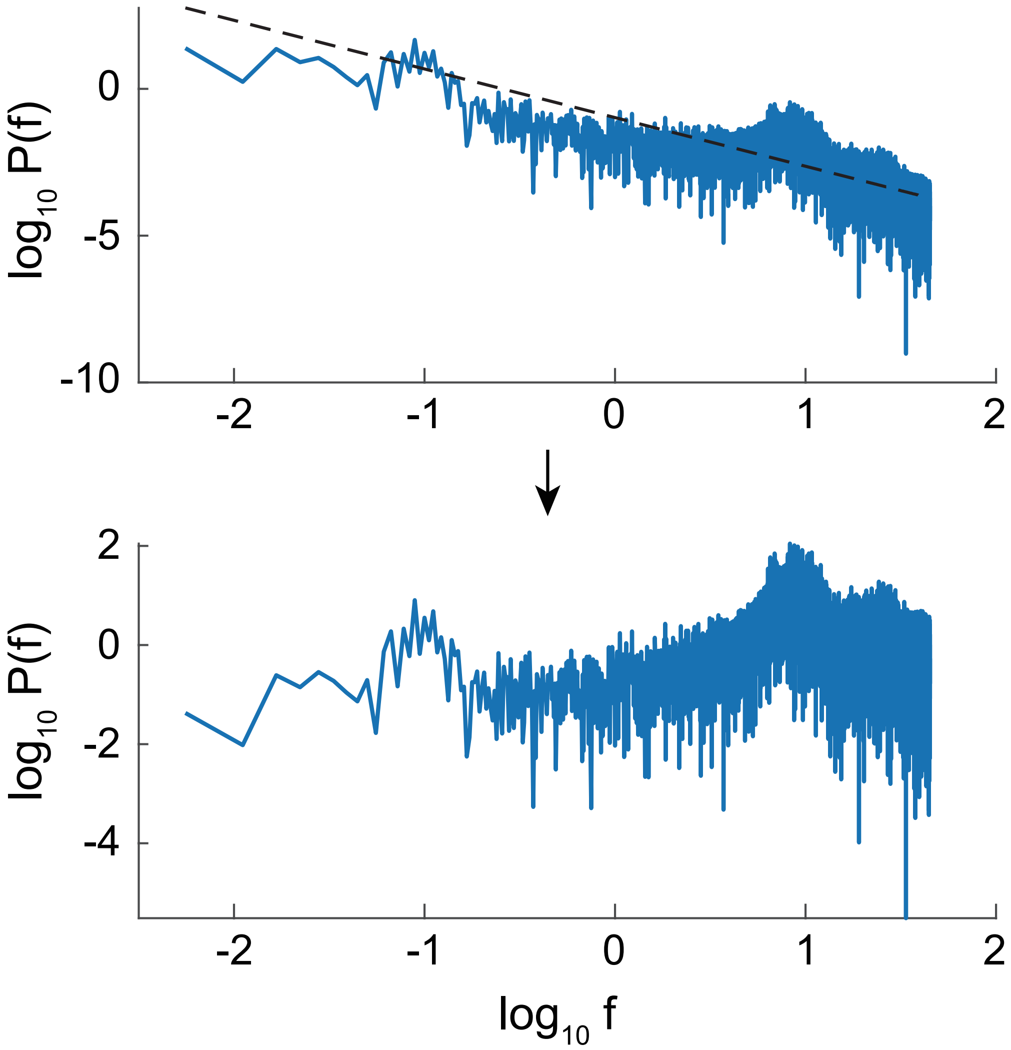


**Fig S2. Estimation of the peak frequency of a fast oscillatory component.** The power spectra of EEG signals were detrended in the double-logarithmic scale with respect to each signal. The detrended spectra were averaged, resulting in a single spectrum from which we estimated the peak frequency (refer to Fig 3E).

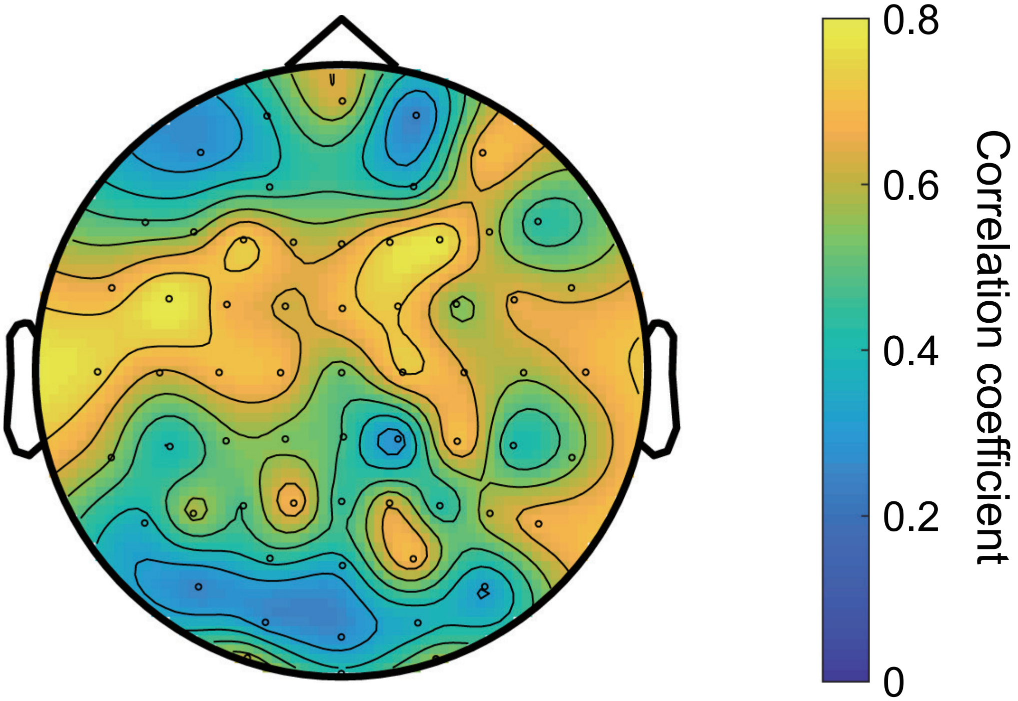


**Fig S3. Correlations between the labeled signals and the time courses of the modulation index (MI) [13].** The MI of the EEG signals was estimated successively over time by a sliding time window with a length of the inverse of the delta-band peak frequency (i.e., one delta wave period). Significant correlations were shown from the scalp sites of more than 50 electrodes.

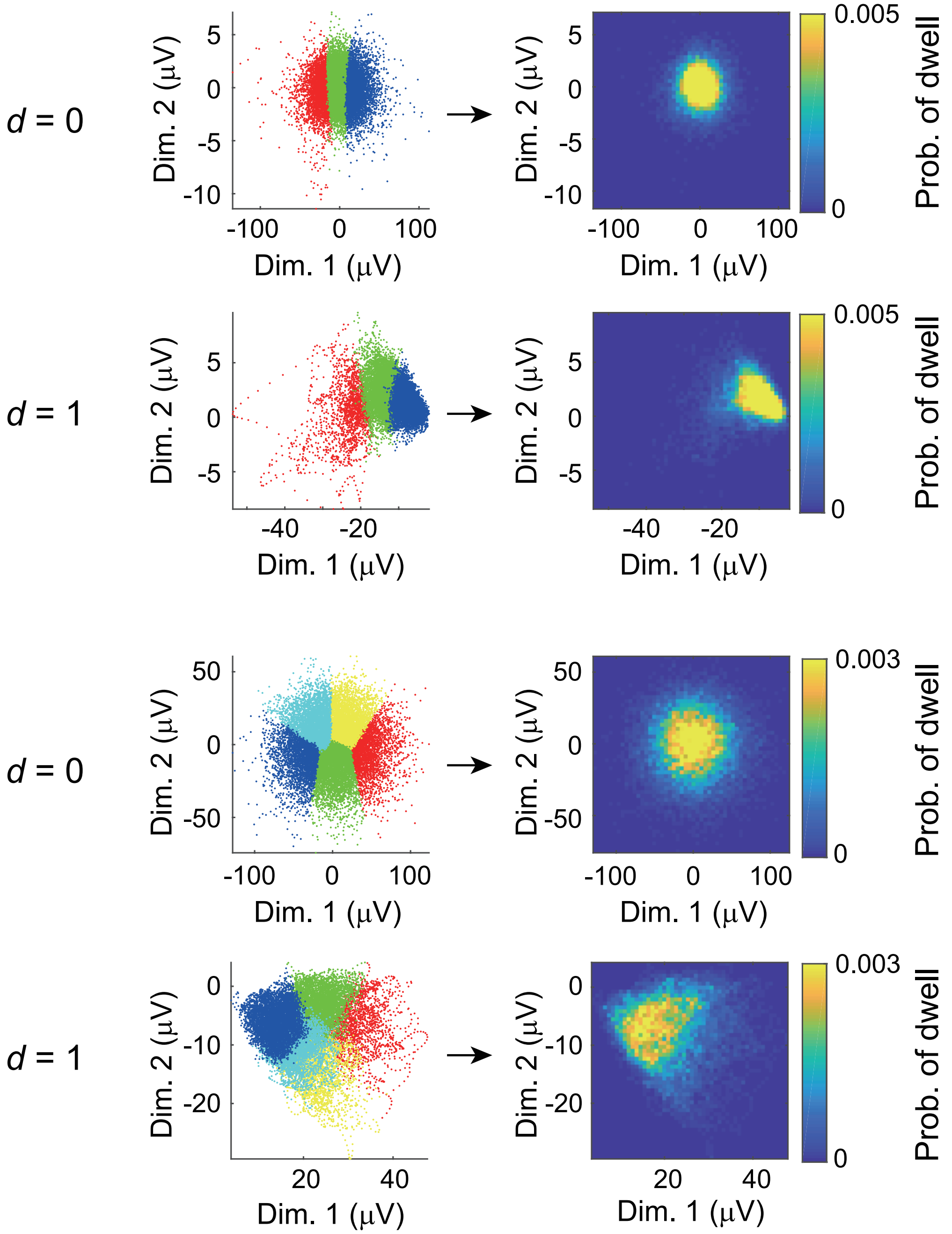


**Fig S4. Trajectories of the experimental delta-alpha PAC dynamics in the $d = 0$ and $d = 1$ conditions.** The surrogate data testing did not reject the null hypothesis $H_0$ in condition $d = 1$ for all the experimental delta-alpha PAC dynamics identified in this study, and many of them were not rejected in the condition of $d = 0$. This Figure corresponds to Figs 4A to 4E, and depicts the case of $K = 3$ estimated from the condition $d = 2$ for comparison purposes.

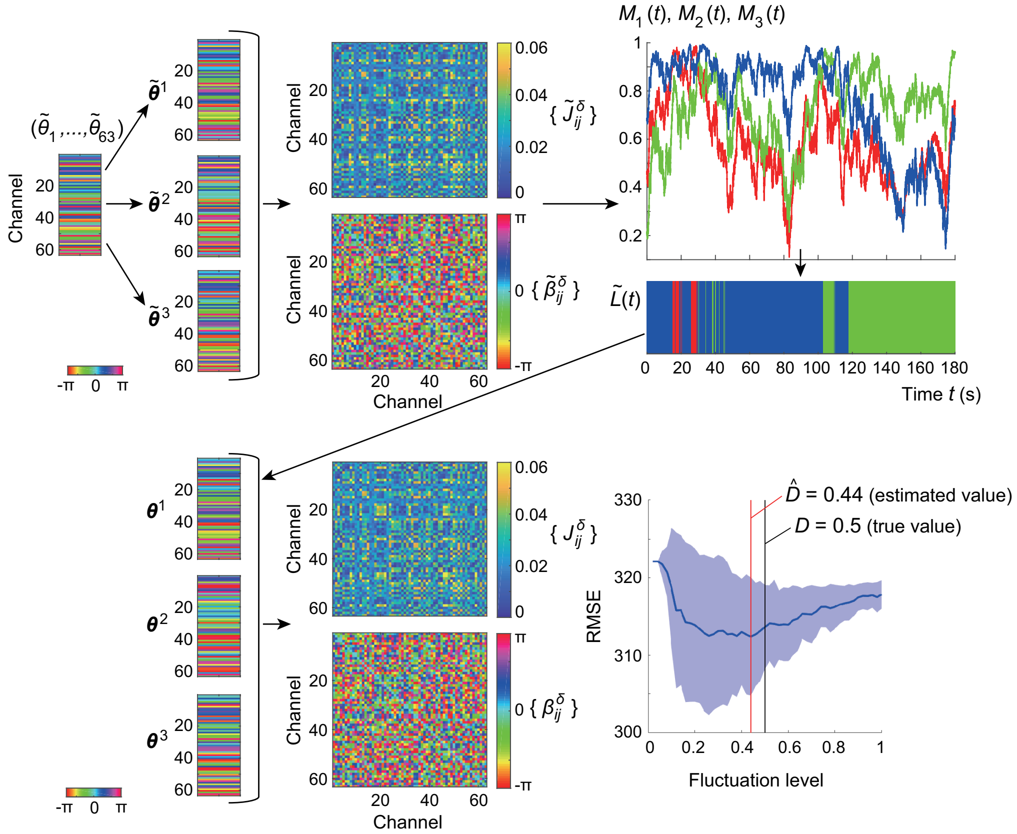


**Fig S5. Validation of the model's parameter estimation.** Artificially generated phase patterns $\{\tilde{\boldsymbol{\theta}}^{\mu}\}$ and the estimated phase patterns $\{\boldsymbol{\theta}^{\mu}\}$ showed high similarities 0.967, 0.881, and 0.962, which were evaluated by $(1/N)|\sum_{j=1}^{N}\exp(i(\tilde{\theta}^{\mu}_j-\theta^{\mu}_j))|$ for $\mu=1,2,3$, respectively. Accordingly, simulated PPC connectivity $\tilde{\boldsymbol{C}}^{\delta}$ and its estimation $\boldsymbol{C}^{\delta}$ showed high cosine similarity (0.979 between the absolute parts and 0.636 between the argument parts). The estimated fluctuation level $\hat{D}$ was 0.44, which was close to the true value of $D=0.5$.

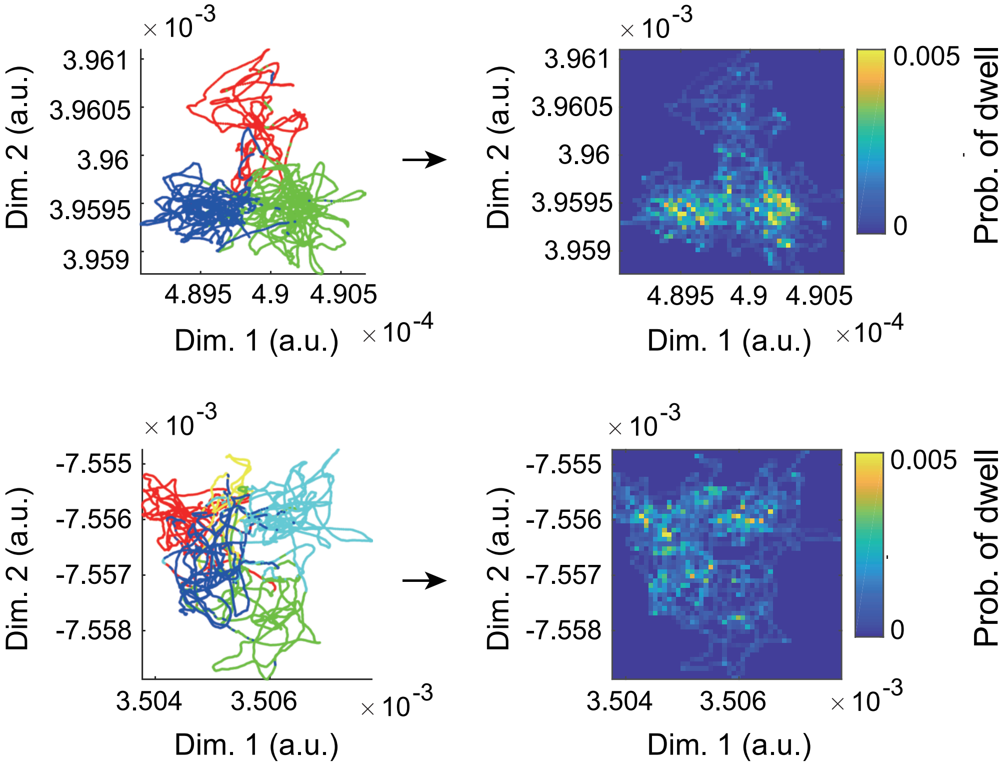


**Fig S6. Trajectories of the modeled delta-alpha PAC dynamics in the condition $d = 1$.** The surrogate data testing did not reject the null hypothesis $H_0$ in condition $d = 1$ for all the modeled delta-alpha PAC dynamics. This Figure corresponds to Figs 8A to 8D, and depicts the case of $K = 3$ for comparison purposes.
